# Innate receptors with high specificity for HLA class I–peptide complexes

**DOI:** 10.1101/2023.02.06.527249

**Authors:** Malcolm J. W. Sim, Paul Brennan, Katherine L. Wahl, Jinghua Lu, Sumati Rajagopalan, Peter D. Sun, Eric O. Long

**Affiliations:** Laboratory of Immunogenetics, National Institute of Allergy and Infectious Diseases, NIH, Rockville, MD 20852

**Author notes:** Correspondence: Malcolm J. W. Sim, Eric O. Long. Current address: Nuffield Department of Medicine, University of Oxford, Headington, United Kingdom (Malcolm J. W. Sim).

## Abstract

Genetic studies associate killer-cell immunoglobulin-like receptors (KIR) and their HLA class I ligands with a variety of human diseases. The basis for these associations, and the relative contribution of inhibitory and activating KIR to NK cell responses are unclear. As KIR binding to HLA-I is peptide-dependent, we performed systematic screens totaling over 3,500 specific interactions to determine the specificity of five KIR for peptides presented by four HLA-C ligands. Inhibitory KIR2DL1 was largely peptide sequence agnostic, binding approximately 60% of hundreds of HLA-peptide complexes tested. Inhibitory KIR2DL2, KIR2DL3, and activating KIR2DS1 and KIR2DS4 bound only 10%, down to 1% of HLA-peptide complexes tested, respectively. Activating KIR2DS1, previously described as weak, had high binding affinity for HLA-C with high peptide sequence specificity. Our data revealed MHC-restricted peptide recognition by germ-line encoded NK receptors and imply that NK cell responses can be shaped by HLA-I bound immunopeptidomes in the context of disease or infection.

## Introduction

Susceptibility and severity of human diseases are influenced by highly variable genes of the immune system, exemplified by the classical class I human leukocyte antigens (HLA-I)^1^. Classical HLA-I molecules (HLA-A, -B and -C) present short peptide antigens that lie in the peptide binding groove (PBG) for immunosurveillance by T cells^2^. Peptides presented by these HLA-I molecules are collectively known as the immunopeptidome and as most HLA-I polymorphism is focused in the PBG, immunopeptidomes are highly diverse and differ between HLA-I molecules^3^. T cell receptors (TCR) detect peptide antigens with exquisite specificity, distinguishing antigens by single amino acid changes and differentiating ‘self’ from ‘non-self’ antigens^2^. However, peptide-specific recognition of HLA molecules is not limited to TCRs, as several studies identified germline-encoded receptors expressed on natural killer (NK) cells that also exhibit peptide-specificity^4–8^. These receptors include members of the killer-cell immunoglobulin-like receptor (KIR) family. In the case of KIR, this peptide-specific recognition contrasts with other innate receptors that bind outside the PBG and do not sense peptide sequence directly, such as CD8, ILT2 and the Ly49 receptors^9, 10, 11^.

KIRs are encoded in a multigene family of activating and inhibitory receptors that play a dominant role in regulating natural killer (NK) cell function^12, 13^. KIRs bind HLA-I towards the C-terminal end of the bound peptide, with positions 7 (p7) and p8 of nonamer peptides being most critical for binding^14–16^. Like their HLA-I ligands, the KIR exhibit considerable allelic polymorphism, in addition to variation in haplotype, copy number, and gene content^12, 17, 18^. Of the 13 KIR genes, six encode HLA-C binding receptors, which are the inhibitory receptors KIR2DL1, KIR2DL2/3, and the activating receptors KIR2DS1, KIR2DS2, KIR2DS4, and KIR2DS5. HLA-C allotypes form two groups, C1 and C2, based on a pair of dimorphic amino acids at positions 77 and 80 that play a central role in determining KIR specificity^19^. C1-HLA-C allotypes (Ser 77, Asn 80) form ligands for the inhibitory receptor allelic pair KIR2DL2/3, while C2-HLA-C allotypes (Asn 77, Lys 80) are ligands for inhibitory receptor KIR2DL1^19, 20^. Activating receptor KIR2DS1, closely related to KIR2DL1 by sequence, also binds C2-HLA-C, but with lower avidity^21, 22^. KIR2DS2 displays no binding to C1-HLA-C on cells but binds C1-HLA-C in the presence of specific peptides, including an epitope conserved in Flaviviruses^20, 23^. KIR2DS4 has an unusual HLA-I specificity, as it binds a subset of C1 and C2-HLA-C allotypes, as well as HLA-A*11^24^. We recently demonstrated that recognition of HLA-C by KIR2DS4 has a high degree of peptide specificity, and that strong ligands include an epitope conserved in bacteria^4^. One KIR2DS5 allotype (KIR2DS5*006) displayed binding similar to that of KIR2DS1, binding all C2-HLA-C allotypes with a lower avidity than KIR2DL1^25^.

The different inhibitory and activating KIR share considerable sequence similarity, bind seemingly analogous ligands, and play a role in regulating NK cell functions. Many studies have linked combinations of specific KIR and HLA-C ligands with human diseases^17, 26, 27^. Homozygosity of both C1-HLA-C and 2DL3 is associated with spontaneous resolution of Hepatitis C Virus (HCV)^28, 29^. Increased risk of pre-eclampsia and low birth weight are associated with pregnancies where the mother carries 2DL1 and the fetus carries its ligand C2-HLA-C^30, 31^. KIR and HLA-C combinations are also associated with susceptibility to autoimmune diseases and the success of hematopoietic stem cell transplantation for treating leukemia^32–34^.

Understanding these associations requires the identification of features that differentiate one KIR-HLA-C interaction from another. Presently, most KIR-HLA-C disease associations are interpreted through the ‘strength of inhibition’ hypothesis^12, 26, 27^. Under this framework, combinations of different KIR haplotypes and presence or absence of specific HLA ligands, generate a hierarchy of NK cell functional states with different predispositions to activation or inhibition. Dominating this hypothesis is the proposition that KIR2DL1-C2 is a stronger interaction than KIR2DL3-C1. Accordingly, 2DL3-C1-HLA-C confers weaker inhibition of NK cell activation, thus allowing NK responses against chronic HCV^28, 29^. Conversely, 2DL1-C2-HLA-C interactions strongly inhibit uterine NK cell activation, thus limiting the NK-dependent transformation of maternal spiral arteries, leading to pre-eclampsia^26, 30^. The evidence for this model relies on the superior binding of recombinant KIR2DL1-Ig fusion proteins (KIR2DL1-Fc) to C2-HLA-C+ cells compared to KIR2DL2-Fc and KIR2DL3-Fc binding to C1-HLA-C+ cells^20^. KIR2DL2 and KIR2DL3 are allotypes and both bind to C1-HLA-C, but KIR2DL2 binds with higher avidity^20, 35^. However, direct affinity measurements of KIR revealed that KIR2DL1, KIR2DL2, and KIR2DL3 have similar affinities for their respective ligands, suggesting that these receptors are not intrinsically stronger or weaker than one another^21, 36–39^. A major difference between these two approaches that measure the strength of KIR-HLA-C interactions is that BIAcore measurements of affinity use recombinant HLA-C, refolded with specific peptides known to bind KIR. In contrast, KIR-Fc based measurements of avidity used HLA-C expressed on cells or cell-derived HLA-C coupled to beads and therefore binding occurred in the presence of a diverse HLA-C bound immunopeptidome^40, 41^.

We have proposed earlier that KIR2DL1 and KIR2DL2/3 exhibit fundamental differences in peptide selectivity, which may provide a better explanation for differences in KIR-Fc binding avidity^14^. Furthermore, we and others showed that activating KIRs have a high degree of peptide specificity and bind peptide ligands conserved in pathogens^4, 23^. Despite the clear contribution of peptide to KIR binding, direct comparisons of KIR peptide specificities have not been performed. Here, we examined the specificity of KIR2DL1, KIR2DL2, KIR2DL3, KIR2DS1, and KIR2DS4 for HLA-C peptide complexes in unprecedented detail using systematic screens to study hundreds of different peptide sequences in the context of both C1 and C2-HLA-C allotypes. Two pairs of C1 and C2 HLA-C with identical sequence, but for amino acids 77 and 80, were chosen to directly compare C1 and C2 allotypes loaded with the same peptides. Our data demonstrate that all these KIR2D are highly peptide sequence specific, except for KIR2DL1 in the context of C2, which is largely sequence agnostic. We categorically show that KIR previously defined as ‘weak’, such as activating KIR, are in fact peptide specific and display HLA-C binding affinities similar to those of inhibitory KIR with optimal peptide ligands. Our data imply that interpretation of disease associations with KIR-HLA-C combinations must consider the degree of peptide specificity and the contribution of immunopeptidomes to NK-target cell interactions.

## Results

### KIR2DL2 and KIR2DL3 are more restricted by peptide sequence than KIR2DL1 for binding to HLA-C

For experiments with recombinant proteins or KIR reporter cell lines, we used the common KIR allotypes KIR2DL1*003 (2DL1), KIR2DL2*001 (2DL2), KIR2DL3*001 (2DL3), KIR2DS1*001 (2DS1) and KIR2DS4*001 (2DS4). We examined KIR binding to peptide libraries loaded onto single HLA-C allotypes expressed in cells deficient in transporter associated with antigen presentation (TAP). To directly compare the specificities of 2DL1, 2DL2, and 2DL3 for the same peptides, we used pairs of HLA-C allotypes that differ only by the C1/C2 dimorphism (positions 77 and 80). The same peptides bind to the C1/C2 pair HLA-C*08:02 and HLA-C*05:01, provided they contain canonical anchor residues^14, 42^. Here, we have also used the C1/C2 pair HLA-C*16:01 and HLA-C*16:02. A comprehensive peptide library based on the ‘self’ 9mer peptide P2 (IIDKSG<u>ST</u>V) was generated where every amino acid combination at position 7 (p7) and p8 of 9mer peptides was included, barring cysteine, totaling 361 peptides (Fig. 1A). Peptide P2 was previously identified from immunopeptidomes of HLA-C*05:01 (C*05) and HLA-C*08:02 (C*08) in multiple studies^14, 40, 41^, and conferred canonical 2DL1 binding to the C2 C*05 and 2DL2 and 2DL3 (2DL2/3) binding to the C1 C*08^14^. All peptides were tested at least twice, with high concordance between experiments (SFig. 1A). Acidic residues (Glu and Asp) at p7 and p8 were incompatible with KIR binding, consistent with previous studies^14, 15, 43^. The 2DL1-C*05 interaction was largely sequence agnostic, with the majority of p7p8 combinations supporting strong KIR binding (Fig. 1A). In sharp contrast, the 2DL2/3-C*08 interaction was more sensitive to peptide sequence, as most p7p8 combinations proved incompatible with binding (Fig 1A, SFig. 1B). 2DL2/3 binding to C*08 required specific p7p8 combinations, dominated by peptides with p8 Ala, Pro, Thr and Ser (Fig. 1A, SFig. 1B). Peptides compatible with 2DL2/3 binding to C*08 were a subset of those that conferred 2DL1 binding to C*05 (Fig. 1B). 2DL2 and 2DL3 displayed near-identical peptide-specificity (Fig. 1C), but 2DL2 binding avidity was stronger than that of 2DL3, consistent with earlier studies^14, 20, 35, 43^.

**Figure 1.**
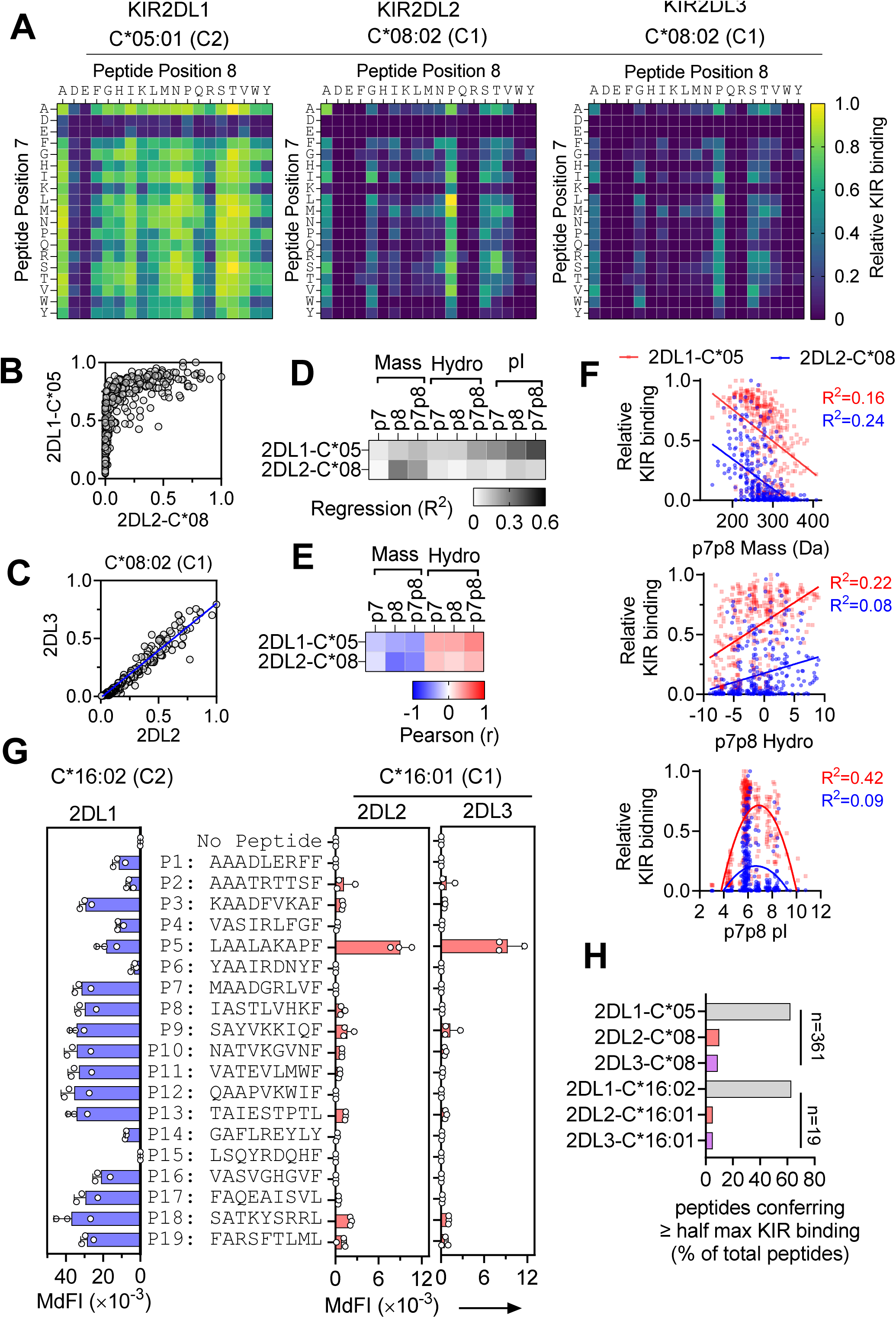
Peptide specificity of KIR2DL1, KIR2DL2, and KIR2DL3 binding to HLA-C. (A) Heatmaps displaying soluble KIR-Fc binding to TAP-deficient cells expressing HLA-C*05:01 or HLA-C*08:02 pre-loaded with a library of peptide P2 (IIDKSGxxV) modified at p7 and p8. For 2DL1-C*05, data are relative to maximum binding with p7p8 ST. For 2DL2-C*08 and 2DL3-C*08 data are relative to maximum 2DL2 binding with p7p8 LP. Mean MdFI (Median fluorescence intensity) of two independent experiments relative to maximum binding as measured by flow cytometry is shown. (B) Correlation of 2DL1-Fc binding to C*05 and 2DL2-Fc binding to C*08, determined from data in (A). (C) Correlation of 2DL2-Fc and 2DL3-Fc binding to C*08 determined from data in (A). (D) R-squared (R^2^) values derived from regression analysis correlating inhibitory KIR-Fc binding with biochemical characteristics at peptide p7, p8 and p7p8 combined. Correlations with mass and hydropathy index (Hydro) were analyzed by linear regression and pI (isoelectric point) with non-linear regression. Mass = amino acid mass (Da), Hydro = hydropathy index, pI = isoelectric point. For mass and hydropathy p7p8 = p7 + p8, for pI p7p8 = (p7+p8)/2. (E) Pearson (r) values from correlations of inhibitory KIR-Fc binding with p7, p8, p7+p8 mass and hydropathy index. (F) Correlation of 2DL1-C*05 and 2DL2-C*08 with p7p8 Mass, hydropathy index and pI. Regression lines and R^2^ values are displayed. (G) 2DL1-Fc binding to C*16:02 (C2), and 2DL2-Fc and 2DL3-Fc binding to C*16:01 (C1) on TAP-deficient cells loaded with 19 C*16:01 self peptides or no peptide was determined by flow cytometry. MdFI from three independent experiments are shown. (H) Proportion of peptides that provide at least half-maximal KIR binding is shown for each of the indicated KIR-HLA-C pairs.

To define the biochemical properties of peptides that bind KIR, we conducted regression analysis of KIR binding correlated with numerical indices of amino acid size (mass), hydrophobicity (hydropathy index)^44^ and isoelectric point (pI)^45^ for amino acids at p7, p8 and p7p8 combined (Fig. 1D-F, SFig. 1C). In general, KIR binding peptides were hydrophobic, and had small side chains and a neutral isoelectric point at p7p8 (Fig. 1D-F, SFig. 1C). The 2DL2-C*08 interaction differed from 2DL1-C*05 by a stronger sensitivity to peptide size, especially at p8, whereas the 2DL1-C*05 interaction favored peptides with hydrophobic amino acids (Fig. 1A, D-F, SFig. 1C).

To explore the generality of our findings in the context of other HLA-C allotypes we generated TAP-deficient cells expressing HLA-C*16:01 (C1) and HLA-C*16:02 (C2) (SFig. 1D,E). As for C*08:02 and C*05:01, this pair of HLA-C allotypes differs only by the two amino acids that define C1/C2 dimorphism. A panel of 19 ‘self’ peptides derived from those previously eluted and sequenced from C*16:01^40^ were synthesized (SFig. 1E). They included every amino acid (except cysteine) at least once at p7 or p8. All peptides stabilized C*16:01 and C*16:02 to a similar extent (SFig. 1E). 2DL1 bound strongly to C*16:02 in the presence of 13/19 peptides (Fig. 1G, SFig. 1F). In sharp contrast, strong 2DL2/3 binding to C*16:01 was observed with only 1 peptide (P5) and much weaker binding was observed with 12/19 peptides (Fig. 1G). Approximately 60% of peptides conferred 2DL1 binding to C*05 or C*16:02 with avidities at least half that obtained with the strongest binding peptide (Fig. 1H). In contrast, only 5-10% of peptides supported 2DL2/3 binding to C*08:02 or C*16:01, respectively, to within 50% of the highest binding peptides. Thus, 2DL1 and 2DL2/3 exhibit very different sensitivities to peptide sequence, with 2DL1 being more sequence agnostic, and 2DL2/3 being more peptide specific.

### Different peptide sequences dictate 2DL2/3 binding to C1 and crossreactive binding to C2 HLA-C

We next explored the capacity of inhibitory KIR to crossreact with non-canonical HLA-C ligands, namely 2DL1 with C1-HLA-C and 2DL2/3 with C2-HLA-C. Consistent with previous studies^14, 35^, 2DL1 displayed exquisite specificity for C2-HLA-C, as only ArgThr at p7p8 conferred 2DL1 binding to C*08 (Fig. 2A, SFig. 2A, B). Crossreactive binding of 2DL2 and 2DL3 with C*05 (C2) was peptide sequence dependent, as it was with canonical binding to C*08 (C1) (Fig. 2A). As observed with C*08 (Fig. 1C), 2DL3 bound C*05 with lower avidity than 2DL2 in the context of the same peptides (Fig. 2B). 2DL3 binding to weaker 2DL2 ligands was very poor when presented by C*05, but not C*08 (Fig. 1C, 2B). Thus, the previously reported lack of C2-HLA-C recognition by 2DL3^+^ NK cells^35, 46^ may be due to a deficiency in 2DL3 to bind weaker 2DL2 ligands. Some p7p8 combinations, such as P2-LP (IIDKSG<u>LP</u>V), were good ligands for 2DL2/3 regardless of the C1/C2 status of the presenting HLA-C (Fig. 2C, D). However, many peptides were detected by 2DL2 selectively in the context of either C1 or C2 (Fig. 2C). For example, P2-QP was a better 2DL2/3 ligand when presented by C*08 (C1), and P2-IQ was a better ligand when presented by C*05 (C2) (Fig. 2C, D). This property was confirmed by cross-reactive 2DL2 binding to C*16:02 (Fig. 2E, F). P5 conferred strong binding to 2DL2 when presented by C*16:01 (C1) but not C*16:02 (C2), whereas P9 and P12 were much better 2DL2 ligands when presented by C*16:02 (Fig. 2E, F). Thus, 2DL2 displays different preferences for peptide sequence when binding the nearly identical C1 or C2 HLA-C allotypes. Finally, the rare crossreactivity of 2DL1 (Fig. 2A)^14^ was confirmed, as none of the 19 peptides loaded onto C*16:01 conferred 2DL1 binding (SFig. 2C).

**Figure 2.**
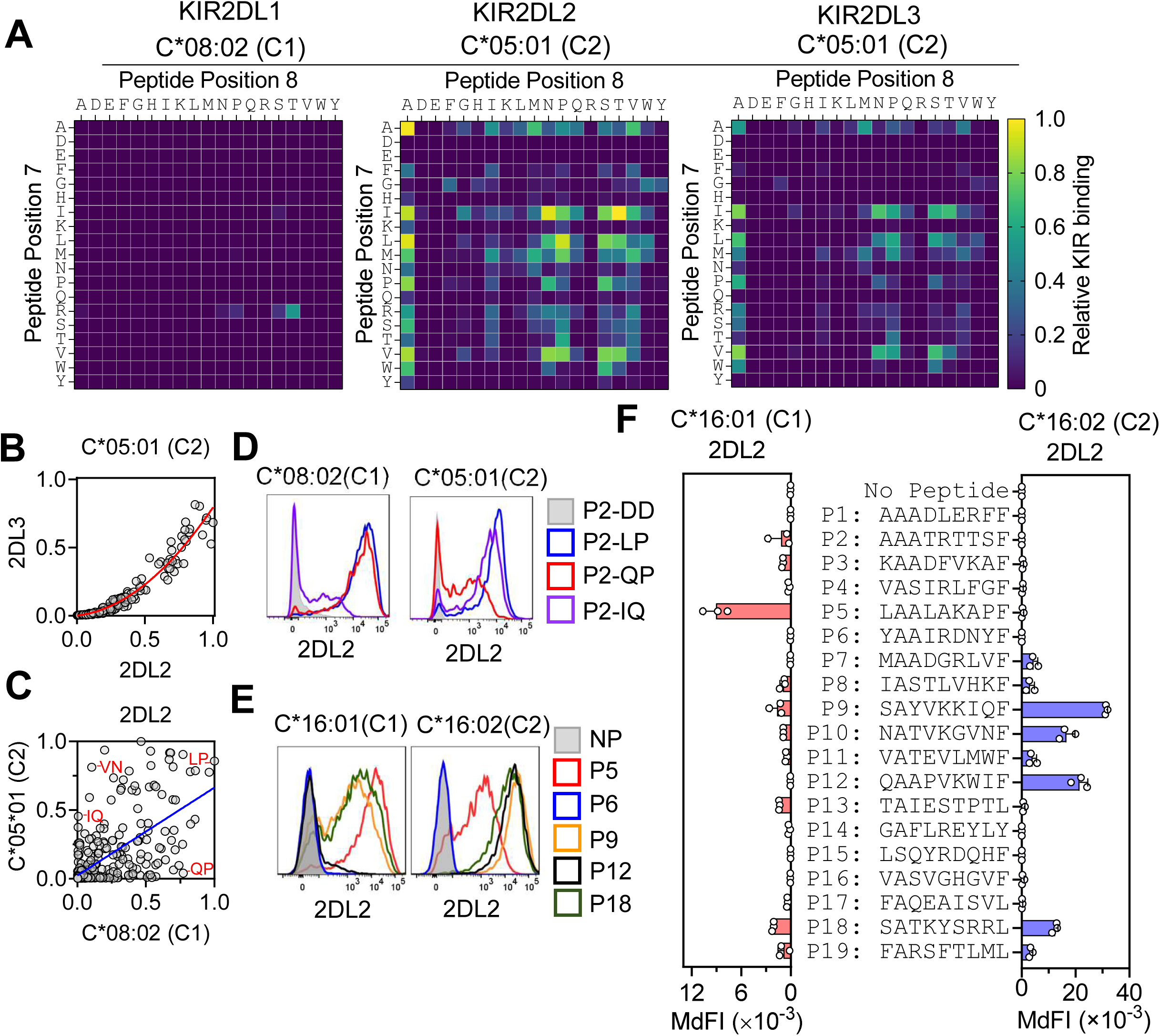
Different peptide sequences dictate KIR2DL2/3 binding to C1 and crossreactive binding to C2 HLA-C. (A) Heatmaps displaying binding of soluble KIR-Fc to TAP-deficient cells expressing HLA-C*05:01 (C2) or HLA-C*08:02 (C1) pre-loaded with a library of peptide P2 (IIDKSGxxV) modified at p7 and p8. Data show 2DL1-Fc binding to C*08:02 relative to maximum binding to C*05:01 with ST, and 2DL2-Fc and 2DL3-Fc binding to C*05:01, relative to maximum 2DL2-Fc binding with IT. KIR-Fc binding was measured by flow cytometry, mean MdFI of two independent experiments are shown. (B) Correlation of 2DL2-Fc and 2DL3-Fc binding to C*05:01, determined from data in (A). (C) Correlation of 2DL2-Fc binding to C*08 and C*05, determined from data in Fig. 1A and 2A. (D) Flow cytometry histograms displaying 2DL2-Fc binding to TAP-deficient C*08 (C1) and C*05 (C2) with indicated peptides. (E) Flow cytometry histograms displaying 2DL2-Fc binding to TAP-deficient C*16:01 (C1) and C*16:02 (C2) pre-loaded with indicated peptides. (F) 2DL2-Fc binding to C*16:01 (C1) and C*16:02 (C2) on TAP-deficient cells loaded with 19 C*16:01 self peptides or no peptide. Data for 2DL2-C*16:01 is the same as in Fig. 1G. Data from three independent experiments are shown.

### High peptide specificity of KIR2DS4 binding to C1 and C2 HLA-C

We recently identified P2-AW (IIDKSG<u>AW</u>V) and similar peptides as functional 2DS4 ligands when bound to C*05^4^. To further dissect the specificity of 2DS4 for HLA-C, we generated a Jurkat reporter cell line expressing 2DS4 and the signaling adaptor DAP12 (Jurkat-2DS4) and screened the P2 p7p8 library. After loading onto C*08 and C*05, 2DS4 recognition of C*08 and C*05 was highly peptide-specific, as only 2/361 peptides bound to C*08 and 7/361 peptides bound to C*05 conferred greater than half-maximal responses (Fig. 3A). 2DS4 ligands were peptides with p8 Trp in specific combinations with p7. For C*08, good responses were seen with MetTrp and ArgTrp, and weaker responses with HisTrp, LeuTrp and IleTrp. Except for HisTrp, these peptides also triggered 2DS4 responses in the context of C*05, but with a different hierarchy (Fig. 3A, B). Additional p7p8 combinations with C*05 triggered 2DS4-specific responses, such as AlaTrp and ValTrp. A p7 side chain was essential as P2-GW did not generate a 2DS4 ligand (Fig. 3A). Thus, 2DS4 binds C1 and C2 HLA-C allotypes, and appeared more peptide-specific when binding C1, as compared to C2-HLA-C (Fig. 3A, B). These p8 Trp containing peptides were validated as 2DS4 ligands by two other assays, KIR-Fc binding and degranulation assays with primary 2DS4^+^ NK cells, after loading onto C*08 and C*05 expressing cells (Fig. 3C-E, SFig. 3A-C). When presented by C*05, many p7p8 combinations are 2DL1 ligands, including those that formed 2DS4 ligands. In contrast, 2DS4 binding peptides presented by C*08 were poor ligand for 2DL2 (Fig 3F). Furthermore, two additional 2DS4 ligands were identified among peptides (P9 and P18) presented by C*16:02 (Fig. 3G). 2DS4 binding to C*16:01 was negative or very weak even in the context of peptides P9 and P18 (SFig. 3D).

**Figure 3.**
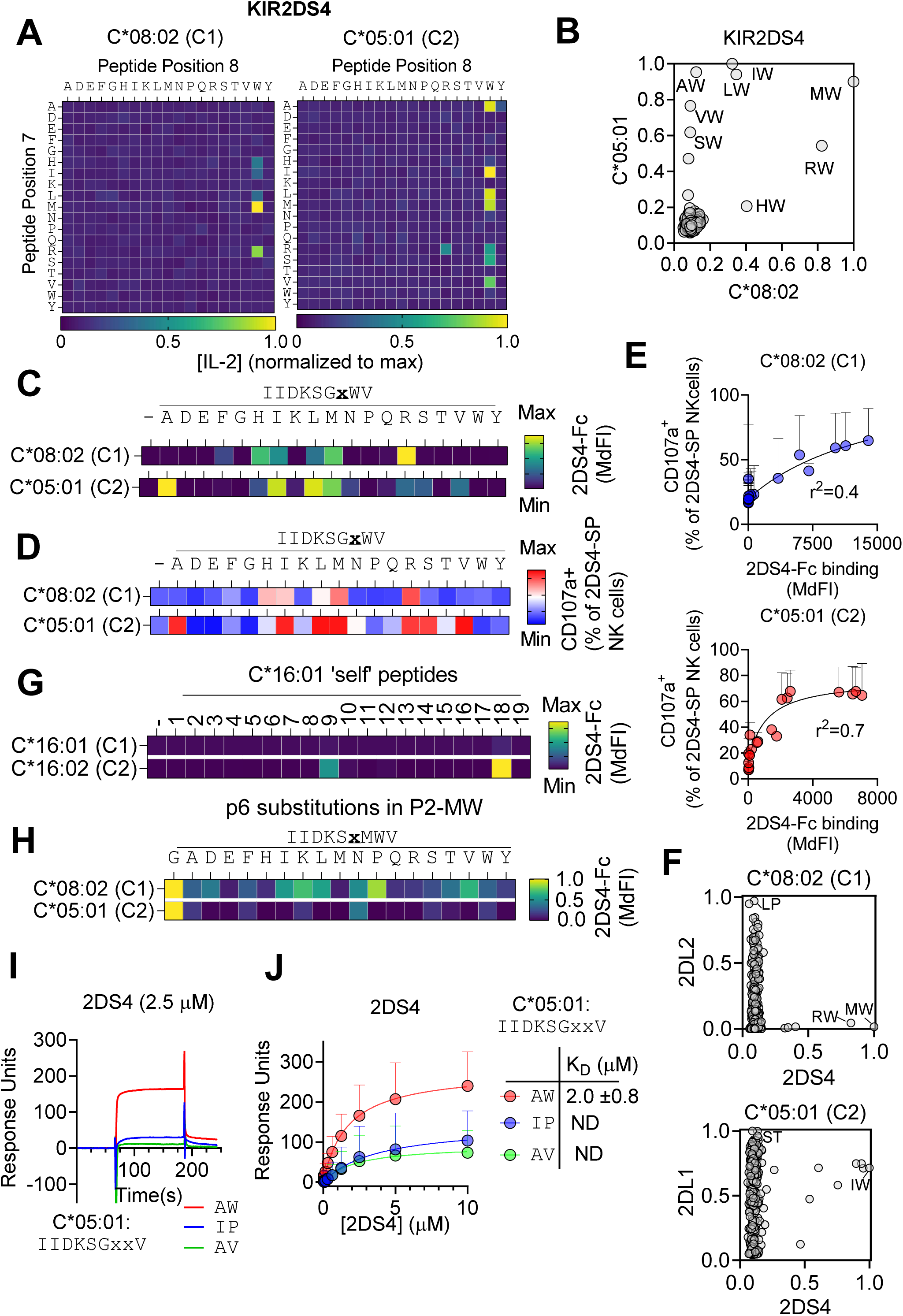
High peptide specificity of KIR2DS4 binding to C1 and C2-HLA-C. (A) Heatmaps displaying IL-2 secretion by 2DS4+ Jurkat T cells in response to C*08 (left) and C*05 (right) pre-loaded with the P2-p7p8 peptide library on TAP-deficient cells. IL-2 secretion was measured by ELISA. Mean of two experiments are shown, normalized to maximal responses with MW for C*08 and IW for C*05. (B) Correlation of Jurkat-2DS4 responses to C*05 and C*08, determined from data in (A). (C) 2DS4-Fc binding to C*05 and C*08 with p8W peptides containing substitutions at p7 of peptide IIDKSGxWV pre-loaded on TAP-deficient cells, as measured by flow cytometry. Mean of three independent experiments is shown. (D) Degranulation of primary resting 2DS4 single positive (2DS4-SP) NK cells in response to C*05 and C*08 cells treated as in (C). Gating for 2DS4-SP NK cells is shown in SFig. 4B. Mean responses of three donors are shown. (E) Correlation of 2DS4-Fc binding shown in (C) and CD107a responses of 2DS4-SP NK cells shown in (D) to C*08 (top) and C*05 (bottom). (F) Correlation of inhibitory and activating KIR recognition of P2 p7p8 library. 2DL2-Fc binding and Jurkat-2DS4 responses to C*08, and 2DL1-Fc and Jurkat-2DS4 responses to C*05. (G) 2DS4-Fc binding to C*16:01 (C1) and C*16:02 (C2) TAP-deficient cells pre-loaded with 19 self peptides (peptide sequences as in 1G). (H) 2DS4-Fc binding to C*08 or C*05 on cells pre-loaded with P2-MW (IIDKSxMWV) and p6 substitutions. Data are mean MdFI from three independent experiments relative to P2-MW with p6 Gly. (I) Representative SPR sensorgrams displaying 2DS4 binding at 2.5 μM to captured C*05 refolded with β2-microglobulin and the peptides P2-AW (IIDKSG<u>AW</u>V), P2-IP (IIDKSG<u>IP</u>V) and P2-AV (IIDKSG<u>AV</u>V). Binding to reference flow cell is subtracted. (J) Equilibrium analysis of surface plasmon resonance (SPR) measurements of 2DS4 binding to captured C*05 refolded with β2-microglobulin and the peptides P2-AW (IIDKSG<u>AW</u>V), P2-IP (IIDKSG<u>IP</u>V) and P2-AV (IIDKSG<u>AV</u>V). Mean responses at each analyte concentration (circles) and standard error between n=5 injections (error bars) are shown. Non-linear regression curve fit of the one-to-one specific binding model (line) are shown. K_D_ value ± standard deviation is shown calculated by modelling steady state kinetics for P2-AW. No K_D_ was determined for 2DS4 binding to P2-IP and P2-AV (ND).

We previously tested 2DS4-Fc binding to twelve C*05 self-peptides that contained p8 Trp and found 1 strong binder (SNDDKNAWF) derived from HECTD11131-1139, demonstrating that 2DS4 ligands can include self-peptides^4^. This peptide has optimal p7p8 residues, however other self-peptides with optimal p7p8 sequences such as IleTrp did not bind 2DS4^4^. As this suggested that other residues influence 2DS4 binding, we examined the contribution of p6 to 2DS4 binding. Substitutions were introduced at p6 in peptide P2-MW (IIDKSxMWV), which is a strong ligand for 2DS4 when presented by C*08 and C*05 (Fig. 3B). In the context of peptide loaded C*08, five p6 substitutions maintained greater than 50% of 2DS4 binding (Pro, Lys, Val, Leu, Ile) (Fig. 3H, SFig. 3G). In contrast, any deviation from p6 Gly decreased 2DS4 binding to C*05, with only p6 Asn conferring more than 25% of binding with p6 Gly (Fig. 3H, SFig. 3G). These data suggest that 2DS4 is more tolerant of variation at p6 when binding C1-HLA-C, but more tolerant of variation at p7 when binding C2-HLA-C (Fig. 3B). The strict preference at p6 for 2DS4 binding to C*05 potentially explains why peptide ISDLDTIWL (UPK3L177-85) did not bind 2DS4 despite optimal p7p8 residues^4^, and further exemplifies the exquisite specificity of 2DS4.

Activating KIR were known as weak receptors, based on binding of KIR-Fc to HLA-C expressed on cells or to HLA-C proteins that had been cleaved from cells and bound to beads^20, 22, 24, 47^. Given our finding that 2DS4 is highly peptide-specific when binding HLA-C, it is possible that a weak avidity to cell-derived HLA-C was due to a paucity of HLA-C peptide complexes compatible with 2DS4 binding. To resolve this question, we determined the solution affinity of 2DS4 for C*05 by surface plasmon resonance (SPR), using recombinant proteins refolded in vitro. 2DS4 bound C*05 refolded with P2-AW (IIDKSG<u>AW</u>V) with low micromolar affinity (K_D_ = 2.0 μM), but not C*05 refolded with P2-IP (IIDKSG<u>IP</u>V) or P2-AV (IIDKSG<u>AV</u>V) (Fig. 3I, J). Thus, 2DS4 is a highly peptide specific receptor that binds C1 and C2 HLA-C with an affinity similar to that of inhibitory KIR (Table 1).

**Table 1.** Affinities (K_D_) of inhibitory and activating KIR2D for peptide:HLA-C. All affinities were determined using recombinant proteins and surface plasmon resonance, except 2DS2 with C*01:02, which used bio-layer interferometry. ^a^Affinity was determined by kinetic analysis of binding curves modelling association and dissociation with KIR concentrations of 2.5, 5 and 10 μM.

| HLA-C | Peptide | KIR |  | K<sub>D</sub> (μM) | Reference |
| --- | --- | --- | --- | --- | --- |
|  |  | Inhibitory | Activating |  |  |
| C*05:01 | IIDKSG**AW**V | 2DL1 |  | 1.1±0.1 | This paper |
|  |  |  | 2DS1 | 1.9±0.2 |  |
|  |  |  | 2DS4 | 2.0±0.8 |  |
| C*05:01 | IIDKSG**AV**V | 2DL1 |  | 1.00±0.1<sup>a</sup> | This paper |
|  |  |  | 2DS1 | 2.6±0.2<sup>a</sup> |  |
|  |  |  | 2DS4 | ND |  |
| C*05:01 | IIDKSG**IP**V | 2DL1 |  | 0.44±0.2<sup>a</sup> | This paper |
|  |  |  | 2DS1 | 2.3±0.8<sup>a</sup> |  |
|  |  |  | 2DS4 | ND |  |
| C*04:01 | QYDDAVYKL | 2DL1 |  | 7.2 | Stewart et al., 2005<br>PNAS |
|  |  |  | 2DS1 | 30.0 |  |
| C*01:02 | LNPSVAATL |  | 2DS2 | 268.7 | Yang et al., 2022<br>Immunology |
| C*06:02 | YQFTGIKKY | 2DL1 |  | 8.9 | Valez-Gomez et al.,<br>2000 Hum. Immunology |
| C*07:02 | RYRPGTVAL | 2DL3 |  | 11.0 |  |
| C*03:04 | GAVDPLLAL | 2DL2 |  | 9.5 | Boyington et al., 2000<br>Nature |
| C*07:02 | RYRPGTVAL | 2DL3 |  | 6.9 | Maenaka et al., 1998,<br>JBC |
| C*07:02 | RYRPGTVAL | 2DL2 |  | 4.5 | Moradi et al., 2021 Nat.<br>Comm. |
| C*07:02 | RYRPGTVAL | 2DL3 |  | 4.2 |  |

### Prediction of novel bacterial ligands for 2DS4

Carrying full-length functional KIR2DS4 is associated with decreased risk of preeclampsia and improved survival in glioblastoma patients, but confers faster progression to AIDS in HIV infected individuals^48–50^. As potential mechanisms for these associations remain unclear, it is critical to develop tools to identify potential 2DS4 epitopes that may contribute to disease. We previously identified a partially conserved 9mer sequence, in the bacterial protein Recombinase A (RecA) that generated a strong 2DS4 ligand when presented by C*05^4^. These peptides were identified by a sequence alignment search of prokaryotic proteomes using P2-AW (IIDKSGAWV). Having defined the 2DS4 binding specificity in much greater detail (Fig. 3), we proceeded to search the UNIPROT database for other potential 2DS4 ligands using the ScanProsite tool^51^ (Fig. 4A). By scanning human, viral and bacterial proteomes with a 9mer search motif that incorporates optimal sequences for binding C*05 (p1-p5 and p9) and 2DS4 (p6-p8) (Fig. 4B), over 500 potential 2DS4 epitopes were identified in bacterial proteomes, as compared to approximately 100 and 50 potential epitopes in the human and viral proteomes, respectively (Fig. 4B). The sequences and source proteins for all predicted 2DS4 epitopes can be found in Supp. Table 1. To validate whether the ScanProsite tool could identify functional 2DS4 epitopes, we focused on bacterial sequences, given their large number. As expected, our search returned many RecA sequences and the remaining were filtered for those conserved in Escherichia coli and other common human pathogens. The Rbba_850-858_ (SLEGPGRWI) and Gudx231-240 (TVDPNGAWL) peptides did not bind 2DS4-Fc or stimulate 2DS4+ NK cells, despite proper loading onto HLA-C (Fig. 4C,D). It is possible that other features besides the simple motif, such as Pro at p4 or p5 in the context of a given sequence, could be incompatible with 2DS4 recognition. The Ehab727-736 peptide (LADNGGAWV) conferred weak 2DS4-Fc binding and elicited an intermediate response by primary 2DS4-SP NK cells, but only when presented by C*05 and not C*08 (Fig, 4C,D). This result is consistent with our data showing that AlaTrp at p7p8 stimulated NK cells only when presented by C*05 (Fig. 3). The TraQ_11-19_ peptide (RLDITGMWV), found in several pathogenic bacterial species (Fig. 4E), was a strong stimulator of 2DS4-positive NK cells when presented by C*08 and C*05 (Fig. 4C,D), again consistent with our data showing that MetTrp at p7p8 was the best combination for binding 2DS4 the context of both C*08 and C*05 (Fig. 3). TraQ is a chaperone-like component of the type IV secretion system, required for bacterial conjugation and horizontal gene transfer^52^. Thus, we have shown that predictions based on experimental data on KIR binding to HLA-C-peptide complexes can be used to identify good ligands for 2DS4, including sequences conserved in bacteria. Detection of pathogens by 2DS4+ NK cells could be relevant to the interpretation of HLA-C and 2DS4 association with disease.

**Figure 4.**
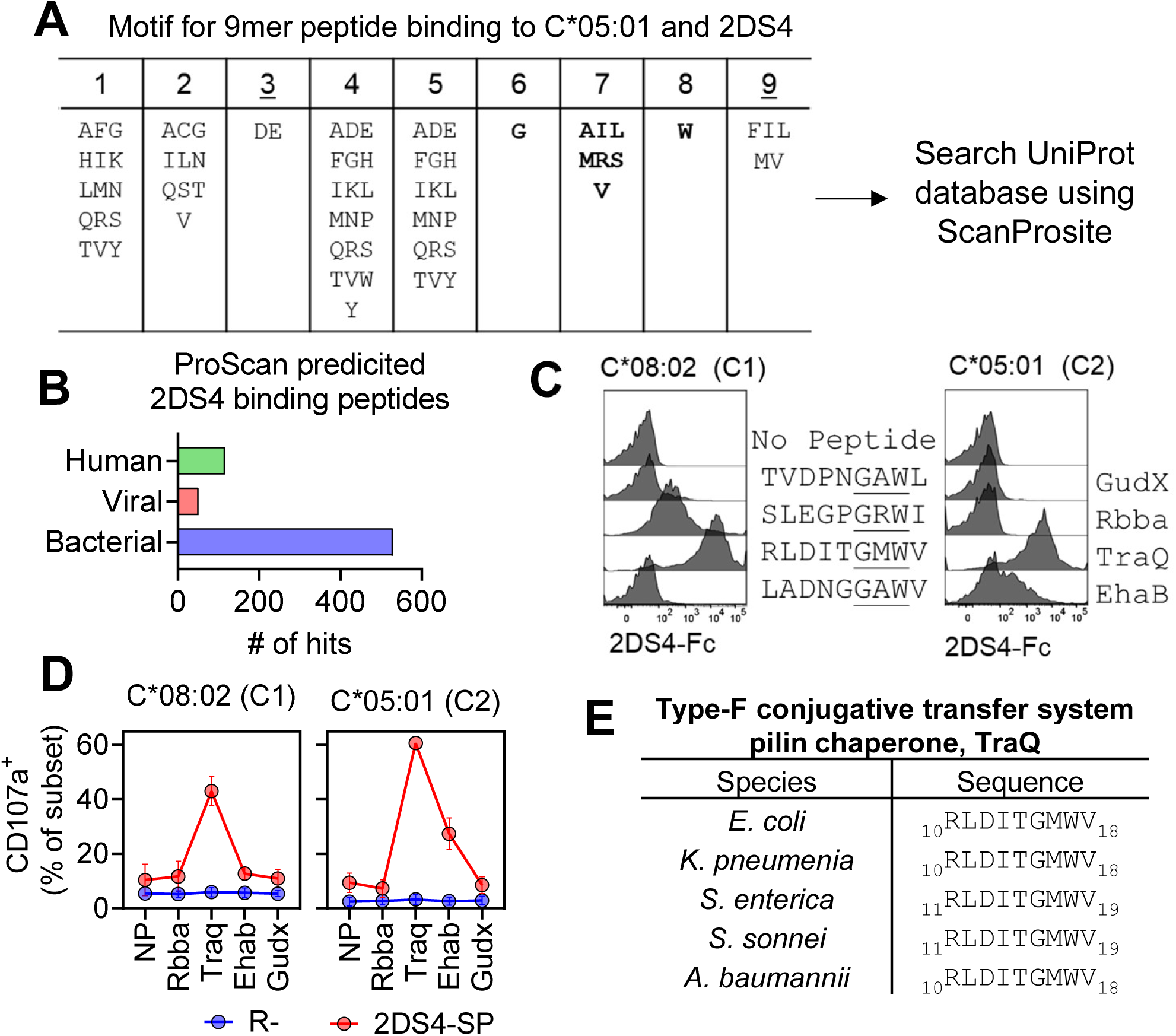
Prediction of KIR2DS4 ligands by sequence homology. (A) Peptide motif used to search UNIPROT database, including amino acids at >1% frequency based on C*05 immunopeptidome. Position 6, p7 and p8 of motif is based on 2DS4 ligands identified in Fig. 4A and 5A. C*05 anchor positions are underlined. 2DS4 binding motif is in bold. C*05 immunopeptidome from Sarkizova et al 2020. (B) Number of potential 2DS4 binding epitopes predicted using motif in (A) and ScanProsite tool (De Castro E et al 2006). (C) Binding of 2DS4-Fc to TAP-deficient C*08 and C*05 cells pre-loaded with indicated bacterial peptides. (D) Degranulation of primary resting 2DS4-SP and Receptor negative (R-) NK cells in response to predicted bacterial 2DS4 ligands pre-loaded on TAP-deficient C*08 or C*05 cells. Gating for R-(2DL1/2/3/S1/S2/S4-, NKG2A-, KIR3DL1-) and 2DS4-SP NK cell populations is shown in SFig. 4B. Data from two NK cell donors are shown. (E) Sequence of TraQ peptide in 5 pathogenic bacterial species.

### High peptide specificity and strong binding of 2DS1 to C2 HLA-C

Unlike 2DS2 and 2DS4, 2DS1 displays measurable binding to cell surface C2-HLA-C and to cell-derived C2-HLA-C bound to beads, with approximately 25-50% of the avidity measured with 2DL1^21, 22, 47^. Studies of approximately 20 peptides presented by C*04:01 and C*06:02 showed that 2DS1 had a preference for peptide sequences similar to that of 2DL1, but bound with lower avidity and affinity^21, 53^. To define the 2DS1 specificity for HLA-C-peptide complexes in greater depth, we used the reporter cell line BW3NG-NFAT-GFP-2DS1, which expresses GFP upon cell surface antibody crosslinking of 2DS1^7^ and detection of peptide-HLA-C complexes (SFig. 4A, B). With the P2 peptide library loaded onto C*05, only 5% (18/361) of peptides conferred greater than half-maximal 2DS1 responses, the highest being P2-VA, P2-LP, P2-IS, P2-IA, and P2-IP (Fig. 5A). Therefore, 2DS1 emerged as a peptide specific receptor, despite 96.9% protein sequence identity with the ectodomain of inhibitory KIR2DL1.

**Figure 5.**
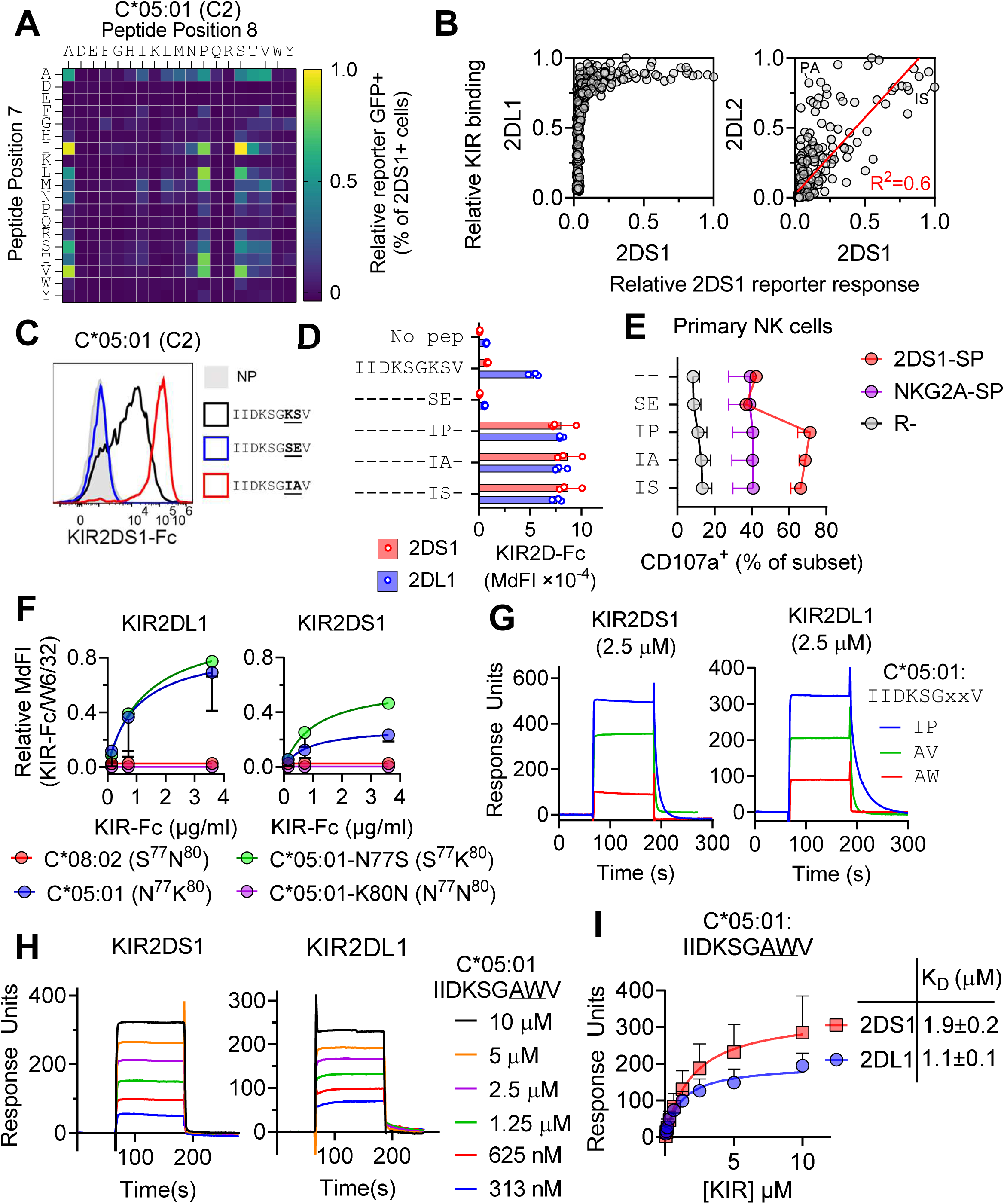
Strong binding and high peptide specificity of KIR2DS1 for C2 HLA-C. (A) Heatmap displaying activation of BWNG-2DS1 reporter cells to TAP-deficient C*05 cells pre-loaded with P2-p7p8 peptide library. Mean responses, relative to maximum response with P2-IS (IIDKSG<u>IS</u>V) from two independent experiments are shown. (B) Correlation of 2DS1 response to C*05 loaded with P2-p7p8 peptide library, as shown in (A), with binding of 2DL1-Fc (data from Fig. 1A) and 2DL2-Fc (data from Fig. 2A). (C) Flow cytometry histograms displaying 2DS1-Fc binding to TAP-deficient C*05 cells pre-loaded with peptides identified in (A) conferring strong (P2-IA; IIDKSGIAV), weak (P2-KS; IIDKSGKSV), and no responses (P2-SE; IIDKSGSEV) to BWNG-2DS1 reporter cells. NP = no peptide. (D) 2DS1-Fc and 2DL1-Fc binding to TAP-deficient C*05 cells pre-loaded with peptides P2-IA, P2-IS and P2-IP, P2-KS and P2-SE. MdFI from three independent experiments are shown. (E) Degranulation of primary resting NK cell subsets in response to C*05 cells pre-loaded with peptides shown in (D). Gating for 2DS1 single positive (2DS1-SP), NKG2A-SP and R-NK cell subsets is shown in SFig. 6B. Data from three NK cell donors are shown (F) 2DL1-Fc and 2DS1-Fc binding to 221-C*05:01 (N77, K80), 221-C*08:02 (S77, N80), 221-C*05:01-K80N (N77, N80) and 221-C*05:01-N77S (S77, K80). KIR-Fc binding is relative to HLA-I expression measured by W6/32. Data shown are mean and standard error from three independent experiments. (G) Representative biacore sensorgrams displaying 2DS1 (left) and 2DL1 (right) binding with 2.5 μM KIR to captured C*05 refolded with peptides P2-IP (IIDKSGIPV), P2-AV, and P2-AW by SPR. Binding to reference flow cell was subtracted. (H) Representative biacore sensorgrams displaying 2DS1 (left) and 2DL1 (right) binding to captured C*05 refolded with peptides P2-AW by SPR. Binding was measured with two-fold dilutions of KIR from 10 μM to 313 nM. Binding to reference flow cell was subtracted. (I) Equilibrium analysis of SPR measurements of 2DL1 and 2DS1 binding to captured C*05 refolded with P2-AW. Mean responses at each analyte concentration and standard error between n=5 injections are shown. Non-linear regression curve fit of the one-to-one specific binding model are shown. Calculated K_D_ value ± standard deviation is shown by modelling steady state kinetics.

2DS1 bound exclusively to the subset of peptides that conferred greater than 75% of maximal 2DL1 binding (Fig. 5B). The top peptides for 2DS1 stimulation were also among those that promoted strong crossreactive binding of 2DL2 with C*05 (Fig. 5B). However, the degree of peptide specificity of 2DS1-induced responses is greater than that of crossreactive 2DL2 binding to the C2 allotype C*05. To confirm results obtained with the 2DS1 reporter cell line, direct binding measurements were performed with soluble KIR-Fc. 2DS1-Fc binding to TAP-deficient, peptide-loaded C*05^+^ cells was indeed as strong as that of 2DL1-Fc in the context of P2-IP, P2-IA, and P2-IS (Fig. 5C, D). Consistent with our screen, 2DS1-Fc binding to P2-KS loaded C*05 cells was much weaker than that of 2DL1-Fc (Fig. 5C, D). As a control, the P2-SE peptide did not support binding of either 2DL1 or 2DS1 to C*05 (Fig. 5C, D).

Functional detection of C*05 by 2DS1 was tested with primary resting NK cells. A gating strategy was used to identify single-positive 2DS1^+^ (2DS1-SP) and NKG2A^+^ (NKG2A-SP) NK cells, as well as NK cells that did not express any inhibitory KIR nor NKG2A (R-) (SFig. 4B). Degranulation was measured after incubation with C*05 cells loaded with peptides that conferred 2DS1-Fc binding. Note that NK cells are normally stimulated by 221 cells, unless 221 expresses an HLA class I for which NK cells have an inhibitory KIR (e.g. C*05 and KIR2DL1). Furthermore, a low response of R-cells was expected, as these NK cells are not licensed. Of the NKG2A-SP NK cells, which benefit from in vivo licensing through inhibitory receptor NKG2A binding to HLA-E, approximately 40%, up from 15%, degranulated (Fig. 5E). Remarkably, loading C*05 on TAP-deficient 221 cells with peptides that provided a ligand for 2DS1 triggered much stronger degranulation, up to 70% of 2DS1-SP cells (Fig. 5E). In the absence of peptide, or after loading P2-SE, degranulation by 2DS1-SP cells was no different from that of NKG2A-SP cells (Fig. 5E, SFig. 4B, C). We conclude that 2DS1 is a strong activating receptor, akin to 2DS4^4^, on primary, resting NK cells and that it endows NK cells with the ability to selectively target cells that have specific peptides presented by C*05. Next, we examined whether the selective recognition of HLA-C-peptides by 2DS1 would apply to another C2 HLA-C allotype. Strong 2DS1-Fc binding to C*16:02 on TAP-deficient 221 cells was detected after loading 3/19 C*16:01 ‘self’ peptides, in contrast to 13/19 peptides that displayed strong binding to 2DL1 (SFig. 4D,E).

The distinct binding properties of 2DS1 and 2DL1 were further examined in the context of the dimorphic residues 77 and 80 in HLA-C that distinguish C1 and C2 allotypes. Selective binding of 2DL1 to C2 HLA-C is dictated mainly by Lys80, which is accommodated by a ’pocket’ in 2DL1^19, 54^. Accordingly, replacing Asn77 with Ser in C*05:01 (mutant S77K80) had no effect on 2DL1-Fc binding (Fig. 5F). Conversely, no binding of 2DL1-Fc occurred with C*08:02 (S77N80) nor with an Asn substitution of Lys80 in C*05:01 (mutant N77N80). 2DS1-Fc binding to these cells was similar to that of 2DL1-Fc but for one striking difference. As with 2DL1-Fc, 2DS1-Fc did not bind to C*08:02 nor to the N77N80 mutant, showing that 2DS1 shares the dependence on Lys80 (Fig. 5F). However, substitution of Asn77 with Ser77 (mutant S77K80) improved 2DS1-Fc binding (Fig. 5F). Ser77 is associated with the presentation of peptides with smaller residues at p8, as compared to Asn77^42^, and smaller p8 residues are favored by 2DL2/3 and 2DS1, which could explain the correlation between peptides favored by 2DS1 and 2DL2 (Fig. 5B).

The reported solution binding affinity (K_D_) of 2DL1 with HLA-C*04:01 refolded with peptide QYDDAVYKL was 7.2 μM^21^. In the same study, 2DS1 displayed a lower affinity for the same HLA-C-peptide complex (K_D_ = 30 μM), suggesting that 2DS1 is a weak receptor. Our work suggests that the peptide may not have been optimal for 2DS1 binding. We therefore determined the solution binding affinity of 2DS1 for C*05 with peptide ligands P2-IP and P2-AV, and the non-stimulating peptide P2-AW. Compared to 2DS4 (Fig. 3I), 2DS1 and 2DL1 displayed stronger affinities, with slower dissociation rates when binding to the strong ligands P2-IP and P2-AV (Fig. 5G, H). For 2DL1 and 2DS1 binding to C*05 refolded with P2-IP and P2-AV, we used kinetic curve fitting to analyze binding curves with KIR concentrations of 2.5 μM, 5 μM and 10 μM. The affinities (K_D_) of 2DS1 for P2-IP and P2-AV were 2.3±0.8 μM and 2.6±0.2 μM respectively, compared to 0.44±0.2 μM and 1.00±0.1 μM, respectively, for 2DL1 (Table 1). The affinity of 2DL1 and 2DS1 for P2-AW was determined by equilibrium binding analysis (steady state) and revealed similar affinities of 1.1±0.1 μM and 1.9±0.2 μM, respectively (Fig. 5I,J). As binding affinities of 2DL1 and 2DS1 were in a similar range, 2DS1 is clearly not a weak receptor. Intriguingly, 2DS1 and 2DS4 bound C*05-P2-AW with similar low micromolar affinities, despite P2-AW not being a functional ligand for 2DS1 (Fig. 4A) and being a potent ligand for 2DS4 (Fig. 3A, D). This suggests that these two activating receptors exhibit different affinity thresholds for receptor activation. Together, these data demonstrate that 2DS1 is not an intrinsically weaker receptor than 2DL1, but a comparably strong binder to HLA-C loaded with peptides that confer high 2DS1 specificity.

### A steep gradient of specificity for peptides presented by HLA-C distinguishes members of the KIR2D family

For a direct comparison of peptide specificity, we took advantage of peptides P9 and P18 presented by C*16:02, which promoted binding of all five KIR studied here. Starting with P9 (SAYVKKIQF) and P18 (SATKYSRRL) backbones, we generated substitution libraries at p8 with fixed p7, and at p7 with fixed p8. Substitutions were made to all other amino acids, except for Cys, Asp and Glu. For 2DL1-Fc binding to C*16:02, only 20% of substitutions reduced binding to less than 25% of the binding to unmodified P9 or P18 (Fig. 6A-D). In contrast, approximately 70%, 55%, and 85% of substitutions reduced binding to that extent for 2DL2/3-Fc, 2DS1-Fc, and 2DS4-Fc, respectively (Fig. 6A-D). Conversely, substitutions that increased binding were less frequent. For the two activating KIR-Fc, some substitutions lead to very large increases in binding (Fig. 6A-D). The Q8S substitution in P9 (SAYVKKI<u>S</u>F) improved 2DS1 binding by approximately 5-fold, and the Q8W substitution in P9 (SAYVKKI<u>W</u>F) improved 2DS4-Fc binding by approximately 90-fold.

**Figure 6.**
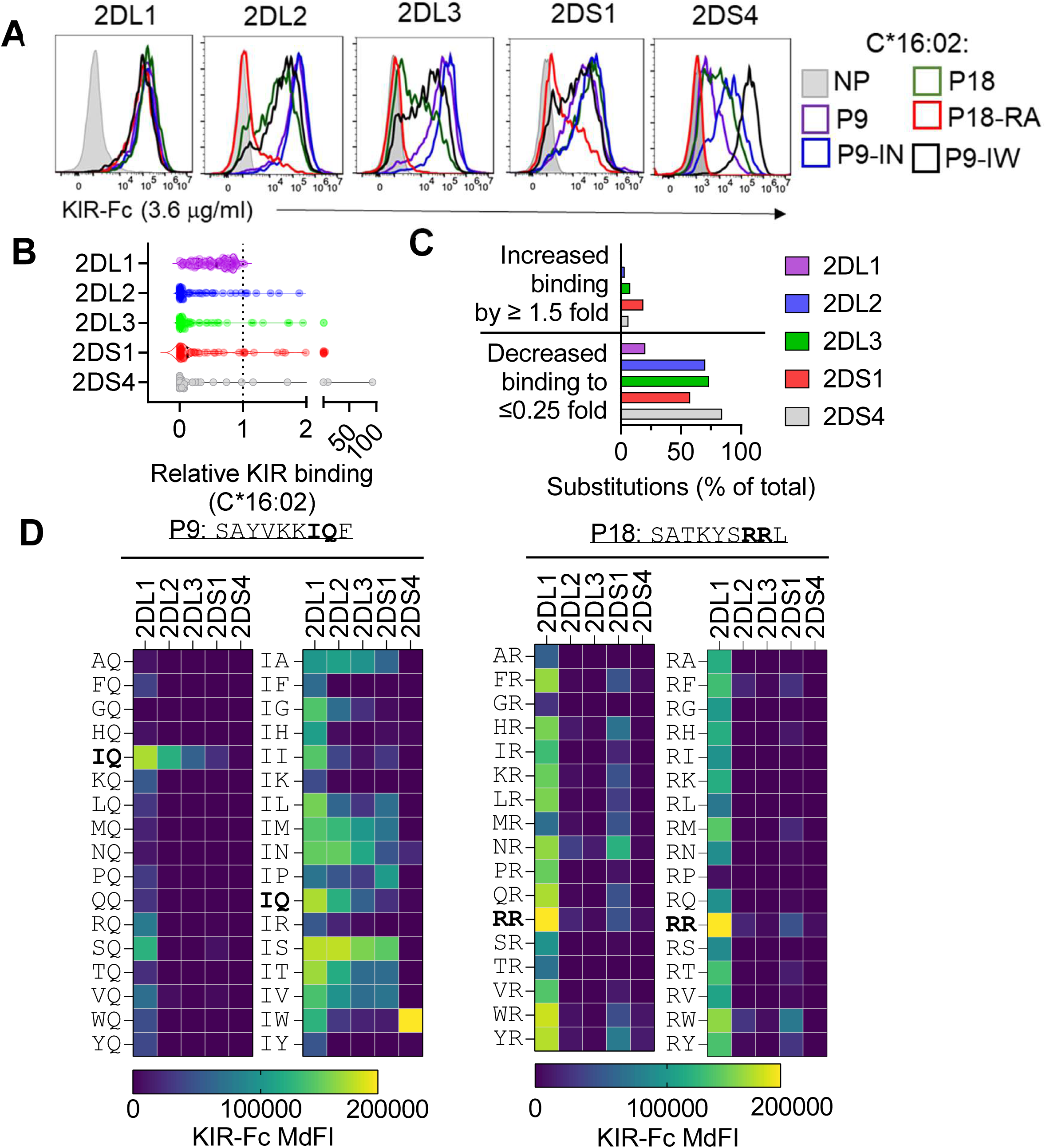
Amino acid substitutions optimize peptide-specific KIR binding to HLA-C*16:02. (A) KIR-Fc (2DL1, 2DL2, 2DL3, 2DS1 and 2DS4) binding to TAP-deficient C*16:02 (C2) cells pre-loaded with P9 (SAYVKKIQF), P18 (SATKYSRRL) and p7p8 substitutions. NP = no peptide, P9-IN = SAYVKK<u>IN</u>F, P9-IW = SAYVKK<u>IW</u>F, P18-RA = SATKYS<u>RA</u>L. (B) Relative KIR binding to C*16:02 loaded with peptides P9 and P18 with substitutions at p7 and p8. (C) Proportion of peptide substitutions at p7 and p8 in P9 and P18 that increase or decrease KIR binding, relative to unmodified peptides. (D) Impact of p7 and p8 substitutions in P9 and P18 on KIR-Fc binding to C*16:02 (C2). Data are mean MdFI from two independent experiments.

The novel 2DS4 binding epitopes were validated functionally using degranulation assays with primary resting NK cells. Peptides that formed better binding sites when loaded onto C*16:02 also stimulated strong degranulation by 2DS4-SP NK cells (SFig. 5). In addition, a few p7p8 substitutions in P9 and P18 improved 2DL2/3 binding to C*16:01 (C1) (SFig. 6) and consistent with our data (Fig. 2), 2DL2/3 binding to canonical (C*16:01) and crossreactive (C*16:02) were optimized by different p7p8 combinations (SFig. 6C, D). These data further demonstrate the peptide specificity of 2DL2/3 in binding HLA-C.

In summary, the 2DL1-C2 interaction was most resistant to peptide sequence substitutions, while other KIR-HLA-C interactions were easily perturbed by sequence changes. In addition, peptide-specific interactions displayed by other KIR could be further optimized with specific amino acids. Thus, 2DL1-C2 differs categorically from other KIR-HLA-C interactions, which exhibit much greater peptide-specificity.

### KIR binding across HLA-C allotypes is dependent on peptide backbone

We next compared KIR binding to different C2 HLA-C allotypes in the context of the same p7p8 peptide combinations: binding to C*05 presenting p7p8 variants of P2 (IIDKSGxxV) compared with KIR binding to C*16:02 presenting p7p8 variants of P9 (SAYVKKxxF) or P18 (SATKYSxxL). Strong positive correlations for all five KIR (2DL1, 2DL2, 2DL3, 2DS1 and 2DS4) were observed with C*05 and C*16:02, when C*16:02 presented peptides with the P9 backbone (Fig. 7A). Notably, p7p8 combinations IS and IW were strong ligands for 2DS1 and 2DS4, respectively, in the context of C*05-P2 and C*16:02-P9. Remarkably, no such correlation occurred between C*05 and C*16:02 that presented peptides on the P18 backbone (Fig. 7B). Similarly, 2DL2 binding to the same p7p8 combinations presented by the C1 allotypes C*08 and C*16:01 showed a stronger correlation when C*16:01 presented peptides on the P9 background than the P18 background (SFig 7). As p7p8 sequence alone cannot explain these correlations, the peptide-specificity of KIR must depend also on structural conformation of p7p8 residues determined by other amino acid positions.

**Figure 7.**
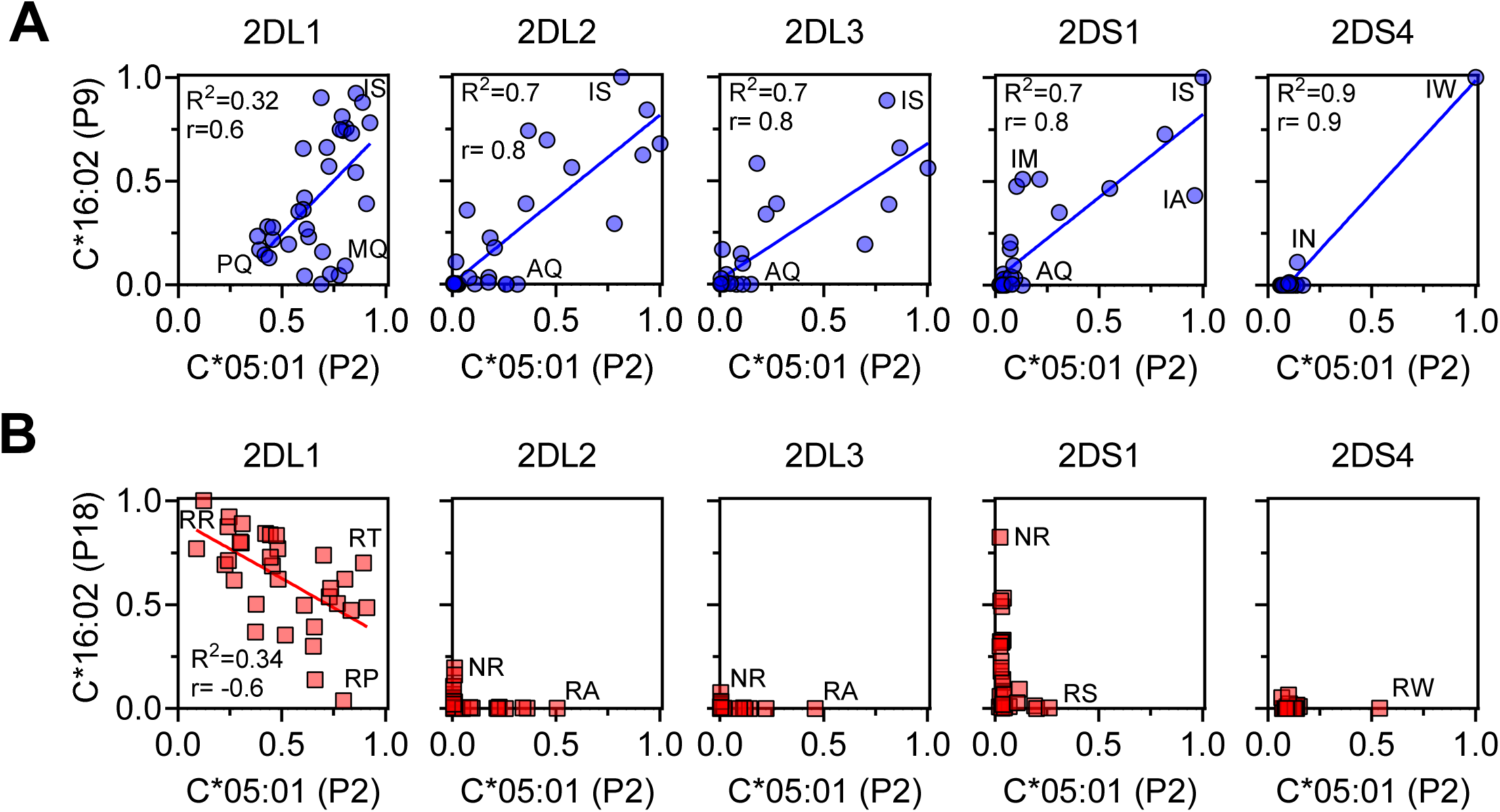
KIR (2DL1, 2DL2, 2DL3, 2DS1, 2DS4) binding to different HLA-C allotypes is dependent on peptide backbone. Correlations of KIR binding to the same p7p8 sequences in the context of peptide P2 (IIDKSGxxV) presented by C*05:01 and peptide P9 (SAYVKKxxF) (A) or P18 (SATKYSxxL) (B), presented by C*16:02. Data are normalized to the maximum value for each peptide backbone.

## Discussion

KIR family members regulate NK cell responses, both positively and negatively, upon recognition of HLA-I ligands. The numerous associations of disease with the presence or absence of specific KIR-HLA-I ligand combinations imply that KIR interaction with HLA-I is not simply generic and that individual KIR members have unique properties. It is therefore crucial to identify salient features that differentiate one KIR-HLA-I interaction from another. We measured over 3,500 different KIR-peptide:HLA-C interactions and uncovered substantial differences in peptide-specificity among KIRs. Notably, 2DL1-C2-HLA-C was unique in being largely peptide agnostic, while four other KIR2D-HLA-C interactions had a sequence dependence ranging from highly selective to exquisitely specific. Furthermore, intrinsic KIR affinity for HLA-C is similar between inhibitory and activating KIR when tested in the presence of optimal peptides, and comparable to the affinity of NK activating receptors, such as 2B4 and NKG2D^55, 56^. Finally, we demonstrate that prediction of KIR ligands from HLA-I-associated peptide sequences is feasible, paving the way for tools that can analyze immunopeptidomes obtained from physiologically relevant tissue sites and identify KIR ligands that may contribute to disease associations.

Differences in peptide-specificity between KIR provide an explanation for the discrepancy observed between KIR binding affinity in solution and binding avidity to HLA-C expressing cells. As cell surface HLA-C presents a diverse immunopeptidome^40, 41^, the frequency of ligands for those KIR that are more peptide-specific is likely to be low, thus resulting in a reduced KIR binding avidity to cells. Thus, our data provide a new lens through which KIR-HLA-C disease associations can be understood, within the framework of the ‘Peptide-selectivity model’^14^. This model proposes that the difference between KIR2DL1-C2 and KIR2DL2/3-C1 is mainly that KIR2DL2/3 are more peptide selective than KIR2DL1. Our current work strongly supports and expands this model, and raises new questions on the contribution of immunopeptidomes to NK cell responses.

Several studies have determined that peptides derived from viruses can be either compatible or incompatible with KIR binding^57–61^. Furthermore, virus infections can significantly alter presentation of host cell-derived peptides^62^, which could alter interactions with KIR. Understanding how the immunopeptidome as a whole contributes to KIR binding, at both steady state and in disease, may reveal the molecular basis for associations with disease. In most cases of KIR associations with disease, the immunopeptidomes of potential NK cell targets, such as fetal extravillous trophoblast (EVT) cells or HCV-infected hepatocytes, are unknown. Our model predicts that in HCV infection, C1-HLA-C infected hepatocytes will present fewer KIR2DL3 ligands, thus conferring weaker KIR2DL3 binding and reduced NK cell inhibition. Conversely, given the promiscuity of KIR2DL1 binding to HLA-I-peptide complexes, immunopeptidomes of C2-HLA-C infected hepatocytes likely provide sufficient KIR2DL1 ligands to maintain NK cell inhibition.

Immunopeptidome studies will have to be paired with tools that accurately predict KIR ligands. As proof-of-concept, we scanned the UniProt database with the KIR2DS4 peptide binding motif identified here, combined with the sequence motif of 9mer peptides bound to HLA-C*05:01, and identified several KIR2DS4 ligands in both human and pathogen protein sequences. Specifically, a sequence conserved in the bacterial protein TraQ was a potent functional ligand for KIR2DS4 when presented by HLA-C*05:01 and the related C1 HLA-C*08:02. The development of more sophisticated computational tools capable of predicting KIR ligands, analogous to MHC peptide binding prediction algorithms^63^, is needed, which will require more input data obtained from screens of similar design to those shown here. The relative contribution of KIR ligand abundance and of ligand affinity to functional NK cell responses has yet to be determined. To what extent signaling by KIR engaged with MHC-I can be modulated by co-activation and co-inhibition receptors, as is the case for TCRs^64^, is not clear. Understanding how NK cells integrate KIR signals with the many other signals they receive from target cells to mount a response could also be a key to parsing KIR-HLA association with disease.

KIR2DS1 is a C2-HLA-C binding receptor with weaker avidity than KIR2DL1^21, 22^. Our data demonstrate that this lower avidity for cell surface HLA-C is due to high peptide-specificity, not an intrinsically lower affinity. Primary resting KIR2DS1-single positive NK cells were stimulated by peptide loaded target cells in a sequence specific manner, indicating that KIR2DS1+ NK cells can contribute to NK cell functional responses when proper ligands are presented. Carrying KIR2DS1 is associated with protection from pre-eclampsia^65^, increased risk of psoriatic arthritis^34^ and better survival in treatment of acute myeloid leukemia (AML) by haemopoietic stem cell transplantation^32^. Future work should define the immunopeptidomes of key cell types proposed to be targets for NK cells in these diseases, such as fetal EVT, keratinocytes, and AML blasts. Cognate KIR2DS1 ligands identified in such immunopeptidomes may provide molecular insight into disease association and expose potential therapeutic avenues.

The broad reactivity of KIR2DS4 with HLA-I allotypes^24^, unusual for a KIR, is best explained by high specificity for a peptide sequence that can be presented by multiple HLA-I allotypes. This could occur through a shared binding motif for KIR2DS4, such as IleTrp at p7p8, which formed a strong ligand on the peptide P2 backbone when presented by C*05, and on the peptide P9 backbone presented by C*16:02. In general, KIR recognition is not limited to p7p8 sequence alone, as the remaining peptide backbone contributed to binding of all KIRs tested, including KIR2DL1. This suggests that KIR recognition imposes both sequence and conformation specific constraints on its ligands, much like the TCR^2, 66^. Functional KIR2DS4 is associated with numerous human diseases, notably with poor outcome in HIV infected individuals^49, 67^. While the mechanism for this association is unknown, it could be due to inflammation triggered by excessive NK cell activation^67^. Immunopeptidomes of HIV-infected and uninfected CD4 T cells may identify KIR2DS4 ligands and provide insight into this disease association.

C2-HLA-C and KIR2DL1, present only in higher primates, are the result of more recent evolution than KIR that bind C1-HLA-C^68^. It is possible that peptide specificity by KIR pre-existed KIR2DL1. As a KIR more resistant to changes in peptide repertoire, KIR2DL1 could provide the advantage of a reliable inhibitory receptor that senses mainly HLA-C expression level. In this respect, KIR2DL1 is like the MHC-I specific Ly49 receptors in mice, which are peptide dependent, but only in so far as a requirement for stable MHC-I, independent of peptide sequence^69, 70^. Activating KIR in chimpanzee have broad and strong MHC-I recognition^47^, indicative of peptide-agnostic recognition, with the exception of KIR2DS4, which is the only activating KIR shared with humans^24^. This suggests that KIR2DS1 and KIR2DS2 evolution in the human lineage generated greater peptide-specificity, perhaps to help combat infections^71^.

KIR2DL2 and KIR2DL3 are alleles of the same gene and exhibit near-identical peptide specificity and similar intrinsic affinities. However, KIR2DL2-Fc exhibits stronger binding avidity than KIR2DL3-Fc when binding to cells or to beads coated with peptide-HLA-C complexes. This is likely due to polymorphism in residues distal to the KIR binding site that are predicted to modify the hinge angle between the D1 and D2 Ig domains^35^. KIR2DL2 was also better than KIR2DL3 at cross-reactive binding with C2-HLA-C, not only due to the lower avidity of KIR2DL3 but also to an apparent deficit of KIR2DL3 in detection of weaker KIR2DL2 ligands^35, 46^. Remarkably, the peptide specificity of KIR2DL2 was different when binding to the same peptides presented by C1 or by C2-HLA-C. In addition to KIR2DL2 and KIR2DL3, allotypes of KIR2DL1 and of KIR2DS1 also exhibit variable binding avidities^72^. Whether this variation in binding avidity is due to differences in peptide-specificity or intrinsic avidity for the same peptide ligands is still unknown.

Inhibitory KIR, along with inhibitory receptor NKG2A, play a dominant role in education (aka licensing) of human NK cells^73–76^. Accordingly, inhibitory receptor interaction with HLA-I maintains NK cells in a state of high responsiveness toward HLA-I-negative target cells. Conversely, interaction of activating KIR with HLA-I can reduce this NK cell responsiveness. KIR2DL1-C2 and KIR2DL3-C1 interactions confer a similar level of education to NK cells^75, 76^, suggesting that despite a greater peptide-specificity of KIR2DL3, the abundance of ligands presented by C1-HLA-C is sufficient to educate KIR2DL3+ NK cells.

KIR2DS1 binding to HLA-C does the opposite of licensing, as it disarms (i.e. reverses education, revokes the license) in C2-HLA-C+ individuals^74, 75^. Despite the high peptide specificity of KIR2DS1, the abundance of ligands on C2-HLA-C for disarming NK cells is sufficient. In contrast, KIR2DS4+ NK cells did not seem to experience disarming^74^. In our experiments, KIR2DS4-SP NK cells were uneducated and hyporesponsive to HLA-I negative target cells, displaying responses similar KIR^-^NKG2A^-^ cells^4^. Altogether, these results suggest that the abundance of specific HLA-C-peptide ligands is insufficient for KIR2DS4 to counter NK cell education. Remarkably, in both cases, 2DS4-SP NK cells^4^ and 2DS1-SP NK cells could overcome the lack of education and trigger NK cell degranulation in response to target cells presenting a specific HLA-C-peptide ligand (Fig. 3D and 4D). In this respect, KIR2DS1 and KIR2DS4 surpass traditional NK activation receptors, which depend on NK cell education for optimal signaling.

Finally, our data cement the KIR in the emerging family of germline-encoded NK cell receptors with specificity for HLA-I-peptide complexes, which include CD94:NKG2C and NKp44^5, 6^. Specific recognition of HCMV peptide-HLA-E complexes by NKG2C+ NK cells stimulates adaptive NK cells in HCMV-infected individuals^6^. Such individuals often expand NK clones with restricted KIR repertoires and expression of activating KIRs^77^. It is possible that recognition of viral peptides by activating KIR occurs also and drives similar clonal, ‘adaptive-like’ NK cell expansions. Indeed, there is evidence that KIR2DS1+ NK cells respond to HCMV-infected fibroblasts in a virus-strain specific manner, indicative of a peptide sequence dependent response^78^. The task ahead is to identify specific peptide ligands for KIR in cells and tissues that may be at the core of the association of KIR-HLA combinations with disease.

## Materials and Methods

### Screening peptide libraries for KIR binding

For peptide libraries, working dilutions of peptide (2 mM) were made in 96 well plates and frozen at -20°C. Using multi-channel pipettes, 10^5^ cells in 190 μl were placed in 96 well plates and incubated with 10 μl of peptide working stock (100 μM final concentration) overnight at 26°C. The following day, cells were stained with KIR-Fc at 4°C for 1hr or mixed with reporter cells at 37°C. For 2DL1-Fc, 2DL2-Fc, 2DL3-Fc and 2DS4-Fc binding to C*05 and C*08, 5×10^4^ 221-C*05-ICP47-GFP and 5×10^4^ 221-C*08-TAP-KO cells were mixed and placed in the same well prior to incubation with peptide. KIR-Fc binding to C*05 (GFP+) and C*08 (GFP-) was determined by KIR-Fc binding to GFP+ and GFP-cells by flow cytometry. For experiments with Jurkat-DAP12-KIR2DS4 cells, 10^5^ TAP deficient C*05 and C*08 cells were incubated with peptide separately overnight at 26°C. The following day, peptide-loaded cells were mixed with 10^5^ Jurkat-KIR2DS4 cells, centrifuged for 3 min at 300 rpm and incubated for 24 hrs at 37°C. Supernatants were collected and frozen. IL-2 concentration in thawed supernatant was measured by ELISA (Biolegend, IL-2 human). For experiments with KIR2DS1 reporter cells, TAP deficient C*05 cells previously loaded with peptide were mixed with 10^5^ KIR2DS1 reporter cells for 6 hrs at 37°C. Cells were washed and stained with anti-KIR2DL1/S1 mAb (EB6-APC, Beckman Coulter) for 30 mins at 4°C. Cells were then analyzed by flow cytometry.

### Peptides

Peptides were synthesized by Genscript (USA). The C*05/C*08 P2 p7p8 library (IIDKSGxxV) consisted of 361 peptides containing all amino acids at p7 and p8 except cysteine and was synthesized at crude purity. The C*16:01/C*16:02 P9 (SAYVKKIQF) and P18 (SATKYSRRL) p7 and p8 libraries consisted of all amino acids except Asp, Glu and Cys. All other peptides were synthesized to 90% purity by Genscript (USA)

### Cells and culture conditions

721.221 cells (221) cells expressing HLA-C*05:01 and TAP inhibitor ICP47 (221-C*05-ICP47) and TAP1 deficient 221 cells expressing HLA-C*08:02 (221-C*08-TAP-KO) were previously described^14, 42^. ICP47 was expressed with GFP via an internal ribosomal entry sequence (IRES). TAP1 deficient 221-C*08:02 cells were generated by expression of HLA-C*08:02 in 221-Cas9 cells followed by transduction with two gRNA targeting TAP1 (Genscript)^42^. BWNG KIR2DS1 reporter cells were a kind gift from Prof. Erik Dissen (University of Oslo, Norway) and have been previously described^7, 79^. BWNG KIR2DS1 reporter cells are derived from BW5147 mouse thymoma cells, where eGFP expression is under control of a 3× NFAT response element promoter. KIR2DS1*001 was expressed as a chimeric fusion protein with mouse CD3ζ cytoplasmic tail, the human CD8 transmembrane region, and the extracellular domains of KIR2DS1 with an N-terminal FLAG tag. Jurkat-DAP12-KIR2DS4 cells were generated by transfection of Jurkat cells with cDNA encoding DAP12 followed by IRES-GFP via nucleofection using Amaxa® Cell Line Nucleofector® Kit V, according to the manufacturer’s instructions. KIR2DS4 was expressed by lentiviral transduction with pcdh encoding KIR2DS4 cDNA. 221 cells were cultured in IMDM plus 10% fetal calf serum (FCS). Jurkat-2DS4 and BWNG-KIR2DS1 reporter cells were cultured in RPMI plus 10% FCS. Peripheral blood samples from healthy US adults were obtained from the NIH Department of Transfusion Medicine under an NIH Institutional Review Board-approved protocol (99-CC-0168) with informed consent. Primary NK cells were isolated by negative selection from peripheral blood mononuclear cells (PBMC) using the NK Cell selection kit (Stem Cell). NK cells were confirmed to be >95% CD56+ CD3-by flow cytometry. Donors for functional experiments were selected by examining KIR expression by flow cytometry. Identification of KIR2DS1 positive NK cells was determined as described using combination KIR2DL1/S1 mAb (EB6, Beckman Coulter) and KIR2DL1 mAb (#143211 R&D), as described^74^. All cells were cultured at 37°C and 5% CO2.

### Recombinant proteins

Recombinant KIR2DL1*003 (#1844-KR-050), KIR2DL2*001 (#3015-KR-050), KIR2DL3*001 (#2014-KR-050) and KIR2DS4*001 (#1847-KR-050) KIR-Ig (KIR-Fc) fusion proteins were purchased from R&D systems. KIR2DS1*001-Fc was custom produced by Sino Biological. KIR-Fc were conjugated to protein-A Alexa 647 (Invitrogen, #P21462) overnight at 1:4 (molar) ratio and used at 3.6 μg/ml unless indicated. Proteins for SPR were generated by refolding from inclusion bodies produced in BL21 (DE3) E. coli (Invitrogen, #EC0114). DNA encoding 2DS1 (3-200) and 2DS4 (3-200) were synthesized and cloned into pET28c by Genscript (USA). Plasmid encoding 2DL1 (1-224) in pET30a was previously described^80^. DNA encoding HLA-C*05:01 (1-278) and β2M (1-99) were synthesized and cloned into pET30a by Genscript and were previously described^42^. Protein expression was induced in bacteria carrying protein expression plasmids grown from single colonies as described^81^. Inclusion bodies were dissolved in 8M urea, 10 mM Tris (pH 8), 150 mM NaCl, 1 mM DTT. For KIRs, 200 mg of dissolved inclusion bodies were refolded by dilution into 1L of refolding buffer containing 400 mM L-Arginine, 100 mM Tris (pH 8), 2 mM EDTA, 0.5 mM oxidized Glutathione, 5 mM reduced Glutathione at 4°C. HLA-C*05:01 peptide complexes were refolded by dissolving HLA-C and β2m inclusion bodies separately before dropwise addition into the same refolding buffer containing 10 mg peptide. Proteins were dialyzed with 10 mM Tris (pH 8) and purified by ion exchanged chromatography with anionic resin (Q columns, Cytvia) followed by size exclusion chromatography with Superdex 200 (GE). Purified proteins were dialyzed against 10 mM Hepes (pH 7.5) 150 mM NaCl prior to SPR studies.

### NK cell degranulation assays

10^5^ TAP deficient 221-HLA-C target cells were incubated overnight with 100 μM peptide at 26°C. The following day, target cells were washed once in RPMI media and mixed with 10^5^ primary NK cells, centrifuged at 300 rpm for 3 mins followed by incubation at 37°C for 2hrs in the presence of BV421 conjugated anti-CD107a mAb (clone H4A3, Biolegend, #328626). Cell mixtures were then washed in PBS and stained with mAbs to CD56 (FITC or BV605, Biolegend, #318304, #318334), NKG2A (PE, clone Z199, Beckman Coulter, IM3291U), KIR2DL1/S1 (PE or APC, clone EB6, Beckman Coulter, A09778, A22332), KIR2DL2/L3/S2 (PE, clone GL183, Beckman Coulter, IM2278U), KIR2DL1 (PE or APC, clone #143211, R&D systems, FAB1844P-100, FAB1844A-100), KIR2DS4 (PE-Cy7, clone JJC11.6, Miltenyi Biotec, 130-130-427) and KIR3DL1/S1 (PE, clone Z27, Beckman Coulter, IM3292). Gating strategies to identify Receptor negative (R-), KIR2DS4 single positive (2DS4-SP) and 2DS1-SP are shown in supplementary figures 3 and 4.

### Surface plasmon resonance (SPR)

SPR experiments were conducted on a BIAcore 3000 instrument and analyzed with BIAevaluation software v4.1 (GE Healthcare). Anti-HLA-I mAb (clone W6/32, Biolegend, # 311402) was conjugated to CM5 chips (Cytvia) at 5000-7000 response units (RU) via amine coupling. HLA-C*05:01-peptide complexes or control HLA-A*03:01-VVVGAGGVGK were captured to 500-700 RU. HLA-C*05:01 was captured on flow cells 2-4 with flow cell 1 left blank or captured with HLA-A*03:01. The analytes were recombinant KIR in 10 mM HEPES (pH 7.5) 150 mM NaCl flown at 50 μl/min. KIR were injected for 2 mins followed by a dissociation time of 5 mins. KIR binding was measured with serial two-fold dilutions of 10 μM to 39 nM. Dissociation constants were determined by modeling steady state analysis, except for 2DL1 and 2DS1 binding to C*05:P2-IP and C*05:P2-AV where dissociation constants were determined by kinetic curve fitting with BIAevaluation software v4.1 using KIR concentrations of 2.5, 5 and 10 μM.

### Protein database search for 2DS4 binding peptides

The UniProt protein database was searched for potential 2DS4 binding peptides using the ScanProsite tool^51^. The motif [AFGHIKLMNQRSTVY]-[ACGILNQSTV]-[DE]-{C}-{CW}-G-[AILMRSV]-W-[FILMV] was entered into ‘Option 2: Submit MOTIFS to scan them against a PROTEIN sequence database’. This search was limited to specific taxa; homo sapiens, viruses and bacteria.

### Flow cytometry

All flow cytometry experiments were carried out on BD Fortessa or Cytoflex (Beckman Coulter) and analyzed using FlowJo software.

## Acknowledgements

We thank Prof. Erik Dissen (University of Oslo, Norway) for kindly providing KIR2DS1 reporter cells. This work was supported by the Intramural Research Program of the NIH, National Institute of Allergy and Infectious Diseases.

## Author contributions

Conceptualization, M.J.W.S., E.O.L., Methodology, M.J.W.S., P.B, J.L.,

Investigation, M.J.W.S., P.B., K. L. W., J.L., S.R.

Writing (original draft), M.J.W.S.

Writing (review and editing), E.O.L. and M.J.W.S. with input from P.D.S., S.R.

Supervision, E.O.L., M.J.W.S, J.L., S.R., and P.D.S.

Funding acquisition, E.O.L. and P.D.S.

## Figure Legends

**Supplementary Figure 1.**
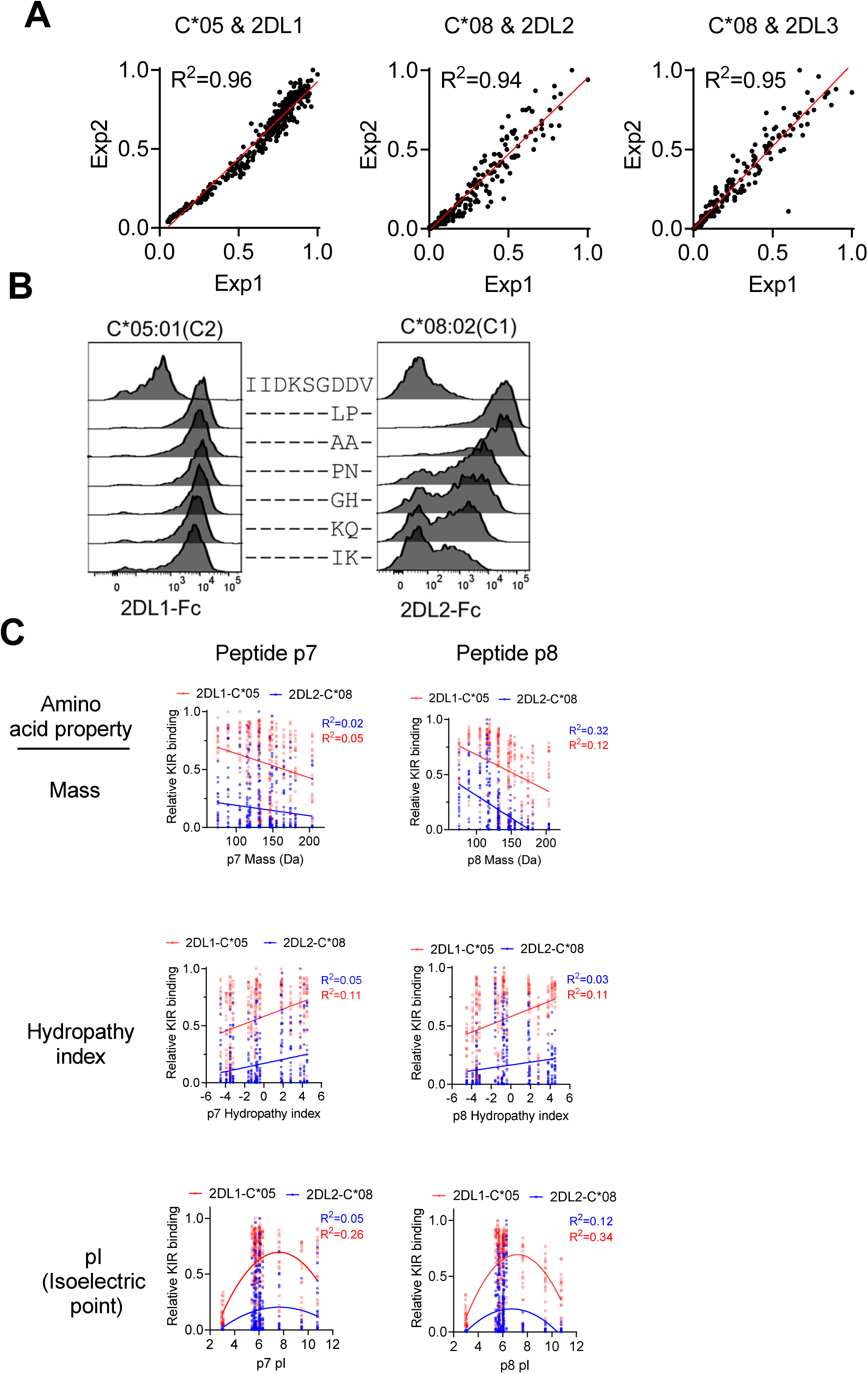
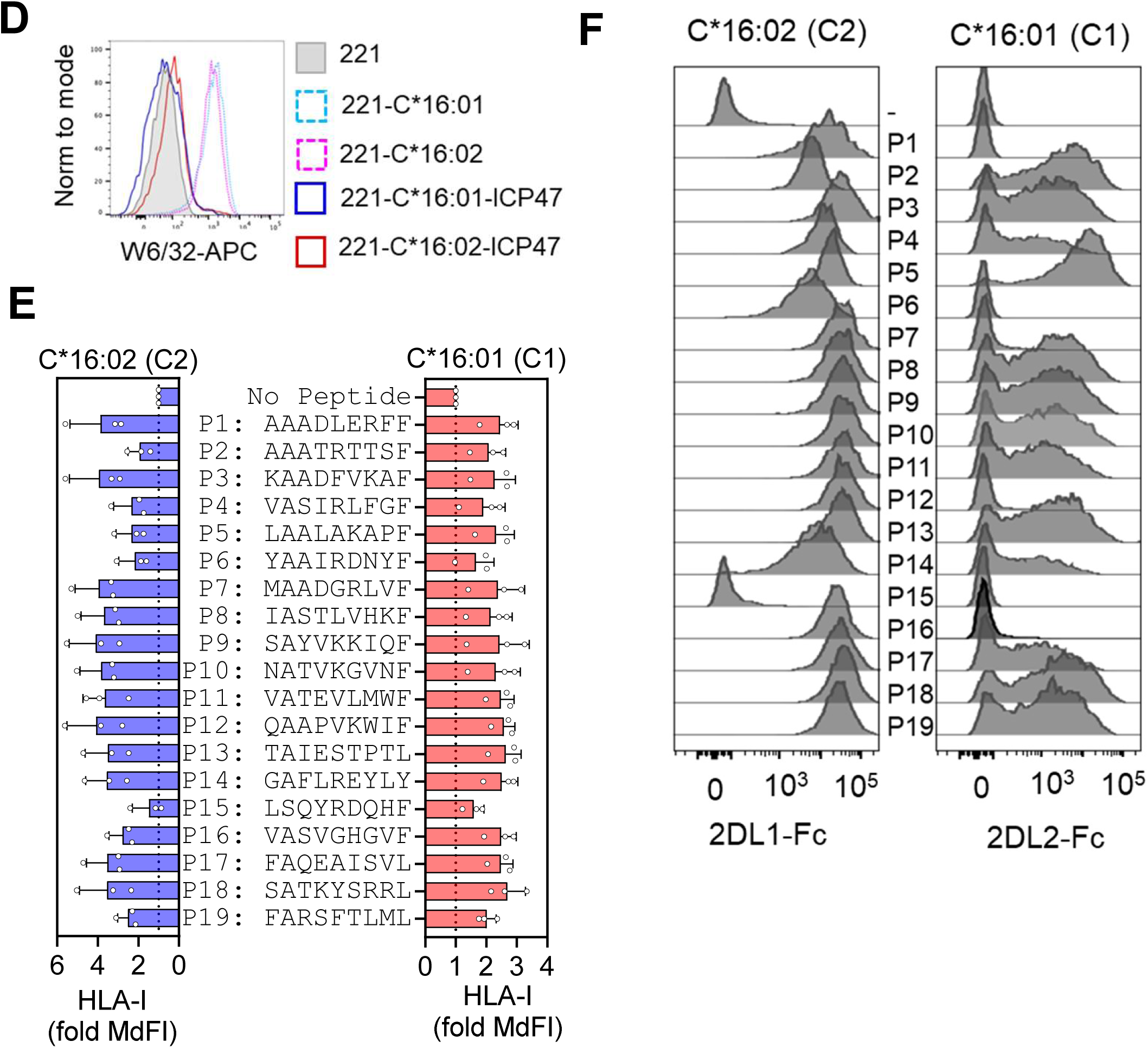
Peptide specificity of KIR2DL1, KIR2DL2 and KIR2DL3 binding to HLA-C. (A) Correlation between repeats of 2DL1-Fc, 2DL2-Fc and 2DL3-Fc binding to C*05 and C*08 in the presence of a 361 peptide library based on p7p8 substitutions in P2 (IIDKSGxxV). As shown in Fig. 1A. (B) Flow cytometry histograms displaying 2DL1-Fc binding to C*05 (C2) and 2DL2-Fc binding to C*08 (C1) loaded with indicated peptides. (C) Correlation and regression analysis of peptide p7 and p8 biochemical properties and KIR2DL1 and KIR2DL2 binding to HLA-C. Linear regression was used for peptide mass and hydropathy index while non-linear regression was used for analysis of pI (isoelectric point). (D) Flow cytometry histograms displaying HLA-I expression on 221, 221-C*16:01, 221-C*16:02, 221-C*16:01-ICP47 and 221-C*16:02-ICP47. (E) Stabilization of HLA-C by indicated peptides loaded on 221-C*16:01-ICP47 (left) and 221-C*16:02-ICP47 (right). (F) Flow cytometry histograms displaying 2DL1-Fc binding to C*16:02 (C2) and 2DL2-Fc binding to C*16:01 (C1) loaded with indicated peptides.

**Supplementary Figure 2.**
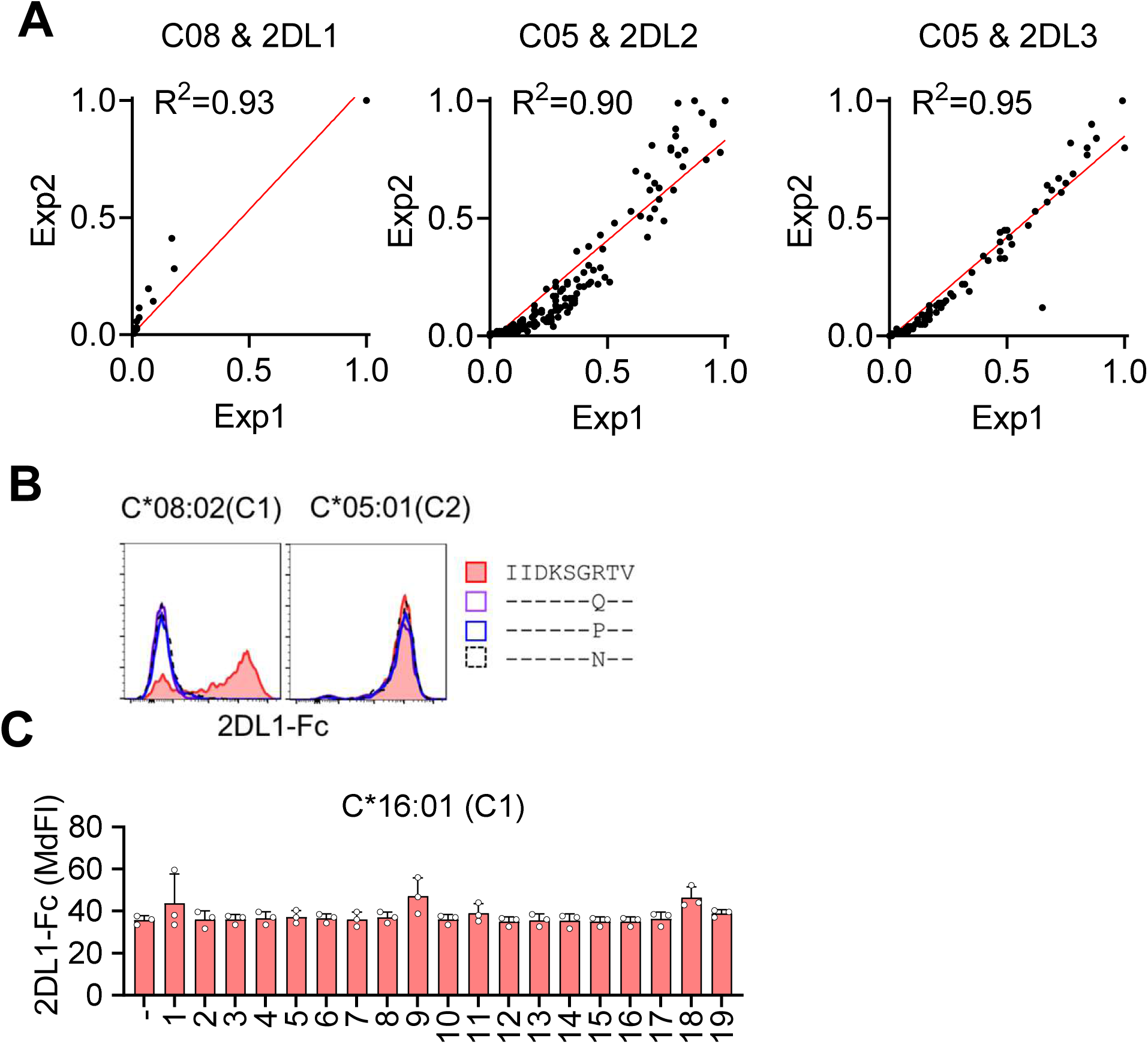
Peptide specificity of crossreactive KIR2DL1, KIR2DL2 and KIR2DL3 binding to HLA-C. (A) Correlation between repeats of 2DL1-Fc, 2DL2-Fc and 2DL3-fc binding to C*08 and C*05 in the presence of a 361 peptide library based on p7p8 substitutions in P2 (IIDKSGxxV). As shown in Fig. 2A. (B) Flow cytometry histograms displaying 2DL1-Fc binding to C*08 (C1) and C*05 (C2) loaded with indicated peptides. (C) 2DL1-Fc binding to C*16:01 pre-loaded with 19 self peptides or no peptide on TAP-deficient cells.

**Supplementary Figure 3.**
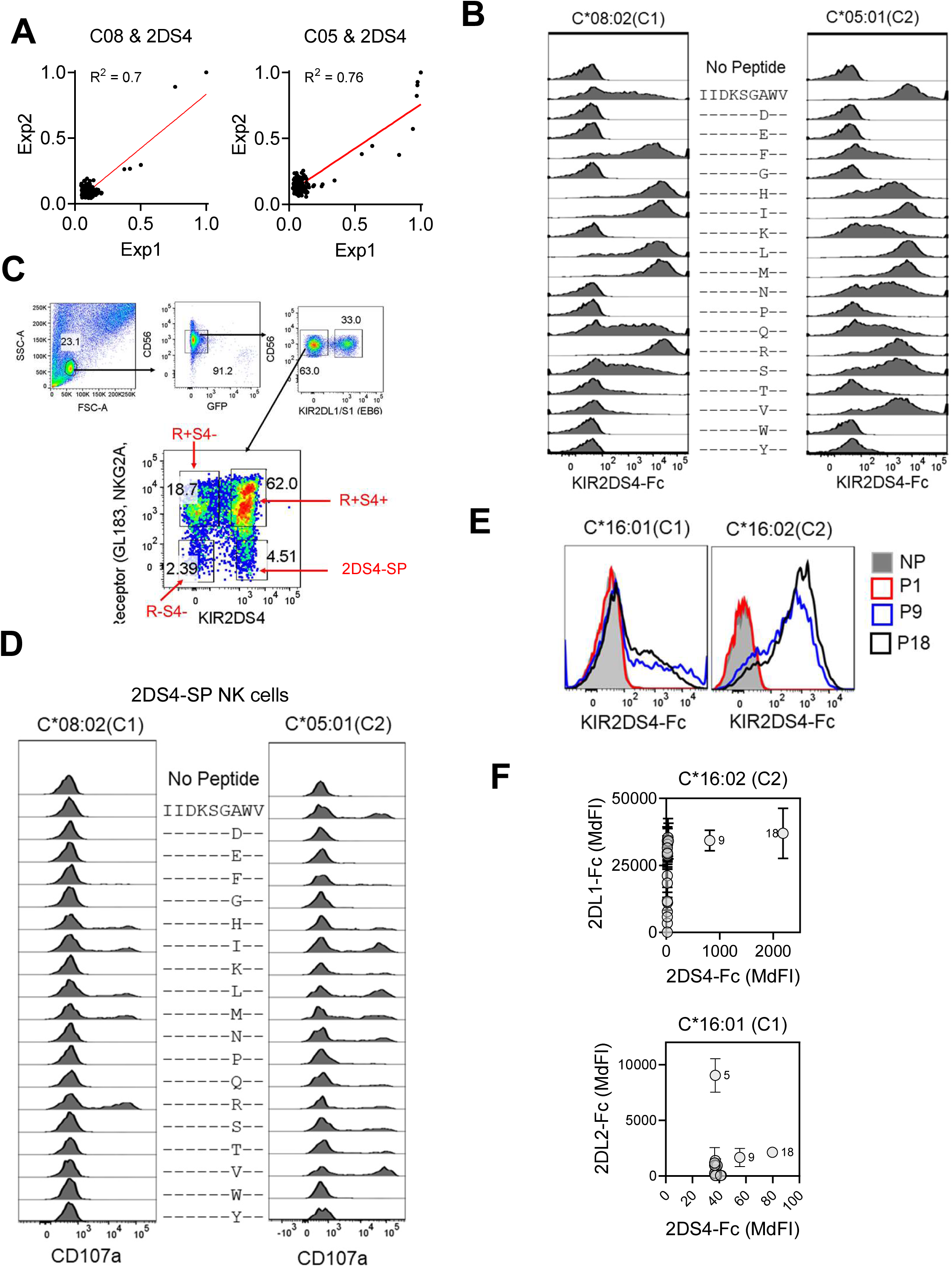
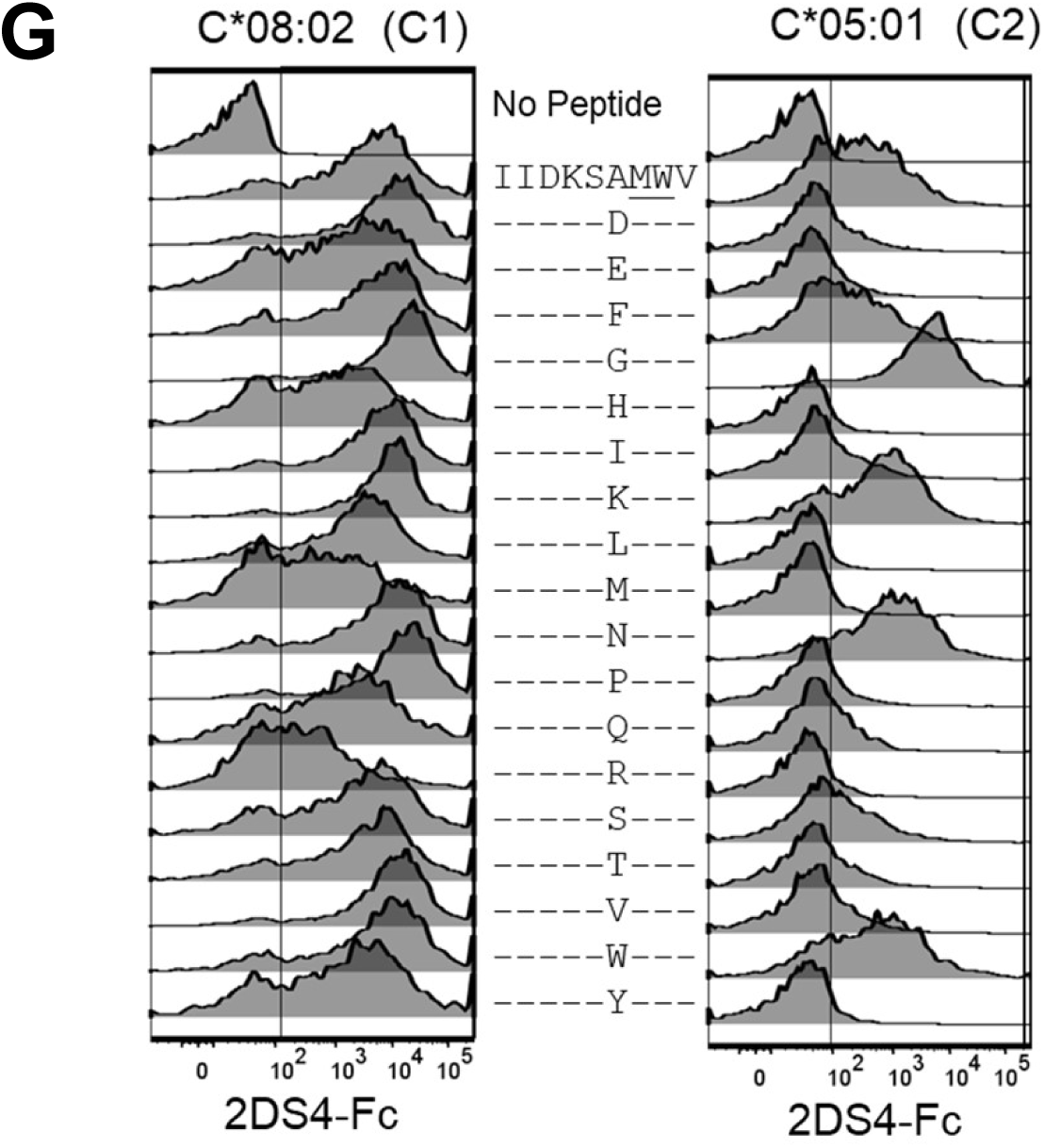
KIR2DS4 is a peptide specific receptor for C1 and C2 HLA-C. (A) Correlation between repeats Jurkat-2DS4 responses to C*08 and C*05 pre-loaded with 361 peptide library based on p7p8 substitutions in P2 (IIDKSGxxV). As shown in Fig. 3A. (B) 2DS4-Fc binding to C*08 (C1, left) and C*05 (C2, right) loaded with p7 variants of IIDKSGxWV. (C) Flow cytometry gating strategy to identify R-S4-(Receptor negative, 2DS4 negative) and 2DS4 single positive (2DS4-SP) subsets. Receptor refers to 2DL2/L3/S2 (mAb GL183), KIR3DL1 (mAb Z70) and NKG2A (mAb Z199). NK cells are identified as CD56+ lymphocytes, GFP-(target cells). NK cells are first gated as 2DL1/S1-(mAb EB6) prior to gating on R and S4. (D) Degranulation of 2DS4-SP NK cells upon mixing with C*08 (left) and C*05 (right) cells loaded with p7 variants of IIDKSGxWV as measured by flow cytometry. (E) 2DS4-Fc binding to C*16:01 (C1, left) and C*16:02 (C2, right) loaded with no peptide (NP), P1 (AAADLERFF), P9 (SAYVKKIQF) and P18 (SATKYSRRL). (F) Correlation of 2DS4-Fc with 2DL1-Fc binding to C*16:02 (top) and of 2DS4-Fc with 2DL2-Fc binding to C*16:01 (bottom). KIR-Fc binding was measured by flow cytometry with peptides pre-loaded on TAP-deficient cells. (G) 2DS4-Fc binding to C*08 (C1, left) and C*05 (C2, right) cells loaded with p6 variants of IIDKSxMWV.

**Supplementary Figure 4.**
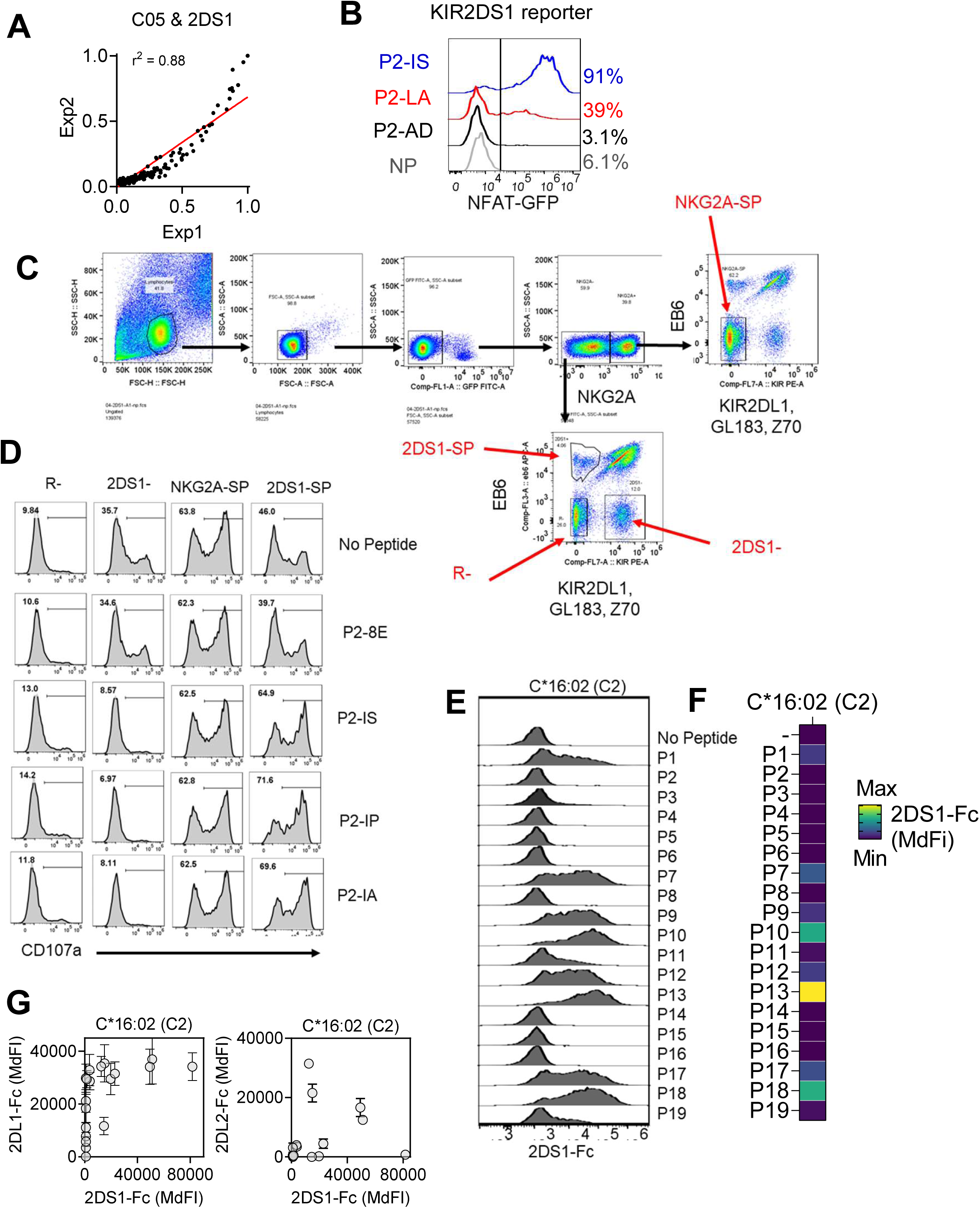
KIR2DS1 is a peptide specific receptor for C2 HLA-C. (A) Correlation between repeat experiments of BWN3G-KIR2DS1-NFAT-GFP responses to C*05 pre-loaded with 361 peptide library based on p7p8 substitutions in P2 (IIDKSGxxV). As shown in Fig. 4A. Data are normalized to maximum GFP+ve cells obtained with p7p8 IS as shown in (B) (B) GFP expression in BWNG-KIR2DS1-NFAT-GFP reporter cells after mixing with peptide loaded C*05 positive cells. NP (no peptide), P2-IS (IIDKSGISV), P2-LA (IIDKSGLAV) and P2-ID (IIDKSGIDV). (C) Flow cytometry gating strategy to identify R-(Receptor negative), KIR2DS1 single positive (2DS1-SP), KIR2DS1 negative (2DS1-) and NKG2A single positive cells (NKG2A-SP). Receptor refers to 2DL2/L3/S2 (GL183), KIR3DL1 (Z70) and NKG2A (Z199). NK cells are identified as CD56+ lymphocytes, GFP-(target cells). (D) Degranulation of NK cell subsets upon mixing with C*05 cells pre-loaded with indicated peptides or no-peptide as measured by flow cytometry. Gating strategy is shown in (C). (E) Flow cytometry histograms displaying 2DS1-Fc binding to C*16:02 (C2) pre-loaded with 19 self peptides on TAP deficient cells. (F) 2DS1-Fc binding to C*16:02 as in (E). Mean MdFI normalized to maximum binding observed with peptide P13 from two independent experiments. (G) Correlation of 2DS1-Fc with 2DL1-Fc, and of 2DS1-Fc with 2DL2-Fc binding to C*16:02 cells. KIR-Fc binding was measured by flow cytometry with peptides pre-loaded on TAP-deficient cells.

**Supplementary Figure 5.**
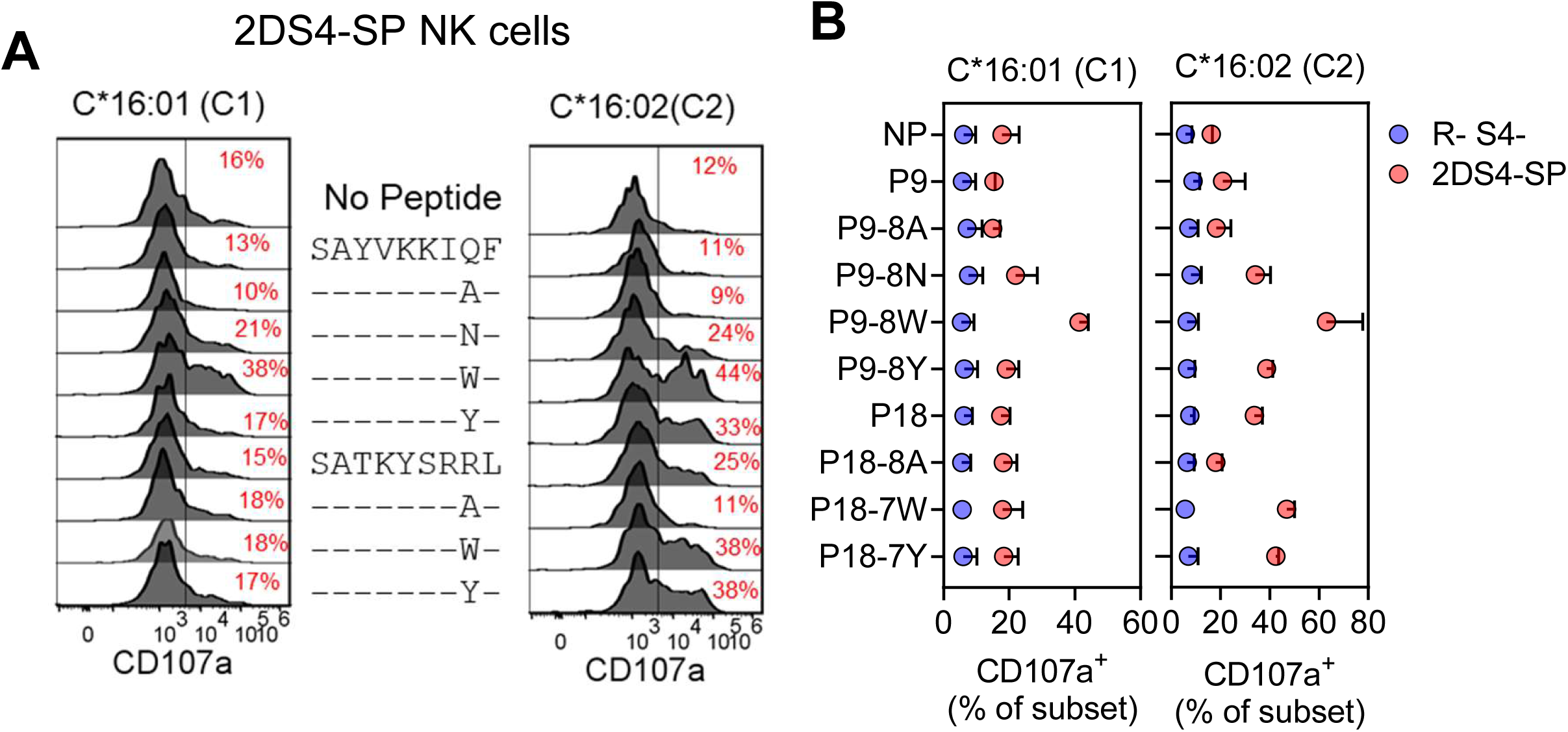
Position 7 and p8 peptide substitutions presented by C*16:02 generate potent KIR2S4 ligands. (A) Flow cytometry histograms displaying expression of CD107a on Receptor negative 2DS4+ (R-S4+) NK cells in response to C*16:01 or C*16:02 cells loaded with indicated p8 variants of P9 (SAYVKKIQF) and P18 (SATKYSRRL). (B) Degranulation of NK cell subsets in response to C*16:01 and C*16:02 cells loaded with peptides shown in (A). R-2DS4- and 2DS4-SP NK subsets were gated as shown in SFig. 3C. Data from two NK cell donors are shown.

**Supplementary Figure 6.**
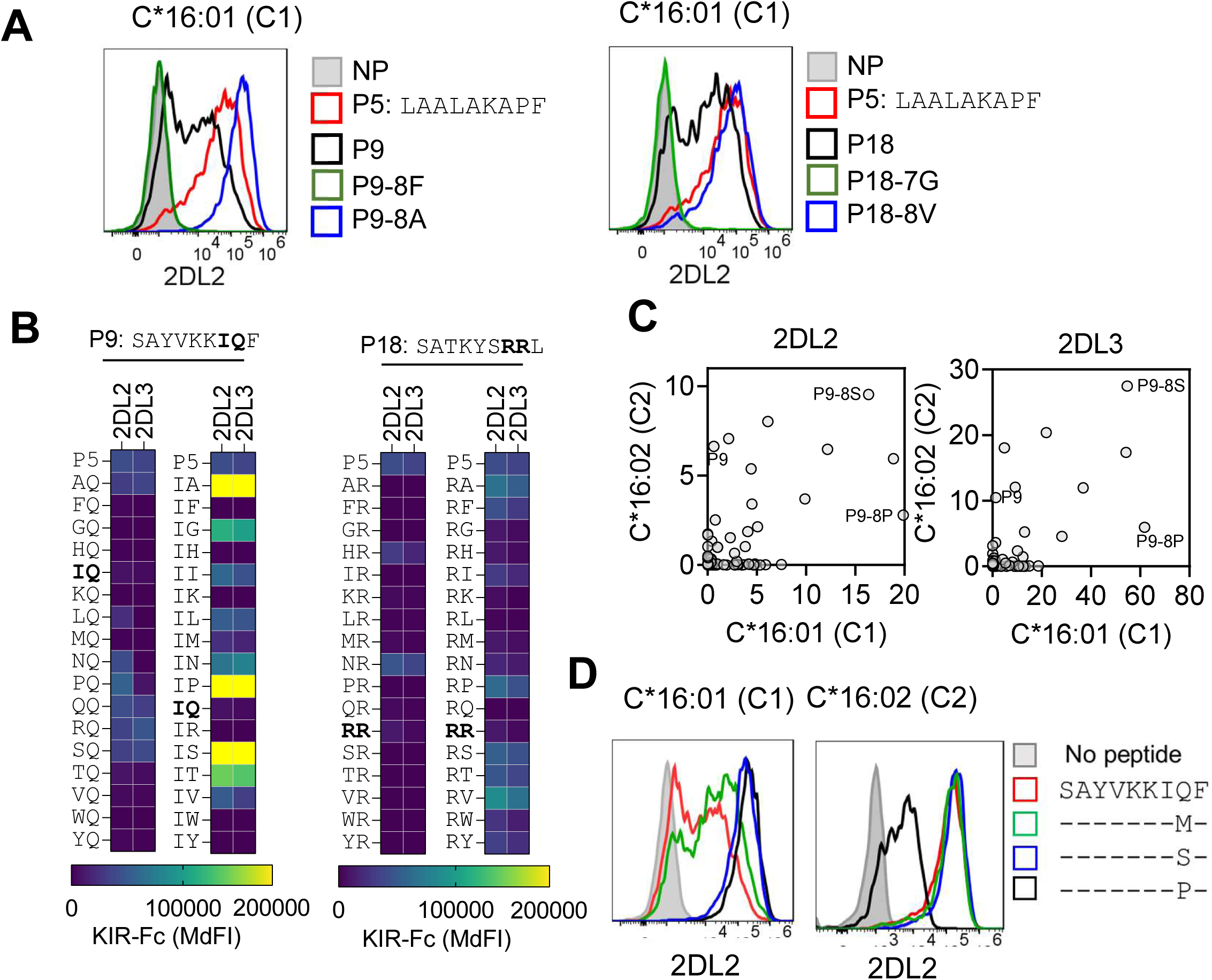
Impact of p7 and p8 substitutions in P9 and P18 on KIR2DL2 and KIR2DL3 binding to C*16:01 (C1). (A) Flow cytometry histograms displaying 2DL2-Fc binding to C*16:01 cells loaded with indicated peptides. NP = no peptide. (B) Relative KIR binding to C*16:01 cells loaded with P9 and P18 with substitutions at p7 and p8. Data are relative to the strong 2DL2-Fc binding to P5, shown in Fig. 1G. (C) Correlation of 2DL2-Fc and 2DL3-Fc binding to P9 and P18 with p7p8 substitutions when preloaded on C*16:01 (C1) and C*16:01 (C2). Data are normalized to binding with P18. (D) Flow cytometry histograms displaying 2DL2-Fc binding to C*16:01 and C*16:02 pre-loaded with indicated peptides on TAP-deficient cells.

**Supplementary Figure 7.**
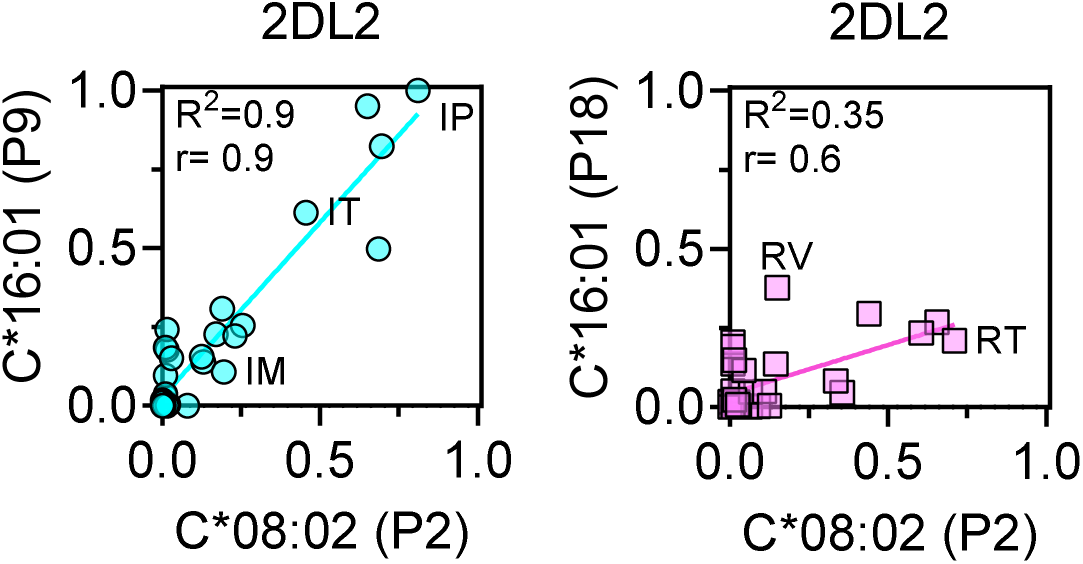
Impact of peptide backbone on 2DL2 binding to C1-HLA-C. Correlation of 2DL2 binding to the same p7p8 sequences in the context of peptide P2 (IIDKSGxxV) presented by C*08:02 and peptide P9 (SAYVKKxxF) (left) or P18 (SATKYSxxL) (right), presented by C*16:01. Data are normalized to the maximum value.

